# Challenging age-related decline in brain function: Evidence from fast neuroimaging of musical sequence recognition

**DOI:** 10.1101/2023.07.13.548815

**Authors:** L. Bonetti, G. Fernández Rubio, M. Lumaca, F. Carlomagno, E. Risgaard Olsen, A. Criscuolo, S.A. Kotz, P. Vuust, E. Brattico, M.L. Kringelbach

## Abstract

Aging is often associated with decline in brain processing power and neural predictive capabilities. To challenge this notion, we used the excellent temporal resolution of magnetoencephalography (MEG) to record the whole-brain activity of 39 older adults (over 60 years old) and 37 young adults (aged 18-25 years) during recognition of previously memorised and novel musical sequences. Our results demonstrate that independent of behavioural measures, older compared to young adults showed increased rapid auditory cortex responses (around 100 and 250 ms after each tone of the sequence) and decreased later responses (around 250 and 350 ms) in hippocampus, ventromedial prefrontal cortex and inferior frontal gyrus. Working memory abilities were associated with stronger brain activity for both young and older adults. Our findings unpick the complexity of the healthy aging brain, showing age-related neural transformations in predictive and memory processes and challenging simplistic notions that non-pathological aging merely diminishes neural predictive capabilities.

## Introduction

Aging is a major omni comprehensive phenomenon which brings new challenges and places large financial burden on society^1, 2^. While most studies have investigated the biological correlates of full-blown disorders such as Alzheimeŕs and other types of dementia^3, 4^, less research has focused on the neural changes associated with normal, non-pathological aging. However, this is crucial to understand the modifications of the brain function and structure across the lifespan and to eventually identify early markers of the age-related neural decline.

Previous research on the neurophysiology of non-pathological aging has predominantly examined age-related changes in resting state brain activity ^5–8^. This research has revealed differences between the spontaneous brain functioning of young versus older adults. For instance, in a magnetoencephalography (MEG) study, Tibon and colleagues ^5^ reported that decreased occurrence of lower-order and increased occurrence of higher-order brain networks were linked to aging. Similarly, combining functional connectivity derived from MEG resting state data with performance in motor learning, Mary and colleagues ^6^ revealed that young and older participants presented differently active neural circuits in resting state after being exposed to motor learning. In another study, Alù and colleagues examined the impact of aging on brain dynamics using electroencephalography (EEG) data and entropy analysis. The findings revealed that older participants had overall higher entropy values across brain regions compared to younger ones ^7^. In another investigation using resting state EEG, the authors found a decrease in occipital delta and posterior cortical alpha rhythms associated with aging ^8^.

Moving away from resting state, a few studies have investigated the impact of aging on automatic brain processes such as the mismatch negativity (MMN), a component of the even-related potential/field (ERP/F) which automatically originates in response to deviant stimuli ^9–15^. For instance, using MEG, Cheng and colleagues ^13^ showed a reduction in the fronto-temporo-parietal activity underlying MMN in older compared to young participants. In another MEG study, the authors revealed that longer peak latencies and smaller amplitudes were found in the MMN of older versus young adults ^14^. Similarly, in an EEG study, Kisley and colleagues showed that older adults presented reduced MMN amplitude at fronto-central sites and decreased sensory gating efficiency compared to younger adults ^15^. Taking together, these findings suggest that aging is associated with declines in automatic central auditory processing of deviant stimuli and with a mild decline of the cognitive ear, possibly related to slow brain atrophy typical of aging ^16^.

The neurophysiology of aging has also focused on memory task-based paradigms ^17–20^. For instance, a few studies suggested that the brain functioning during long-term recognition is impaired in older compared to young adults. Gajewski and Falkenstein ^17^ revealed decreased and delayed ERP components (e.g. N200, P300a and P300b) in older adults when they performed a two-back memory task. Along this line, using EEG, Vaden and colleagues ^18^ showed that in a suppression of visual information task, the correct performances were associated with a robust modulation of alpha power only in young but not in older adults. Similarly, Federmeier and colleagues ^19^ demonstrated that older compared to young adults had a reduced neural efficiency when recognising familiar words.

Previous research has also investigated the impact of aging on short-term recognition of information, showing altered brain functioning in older compared to young adults. In a classic MEG study, Babiloni and colleagues ^20^ used two delayed response tasks, reporting altered alpha event-related desynchronisation (ERD) associated with aging. In a recent EEG study, Costa and colleagues ^21^ investigated the age-related differences in the neural activity during short-term recognition of musical sequences. They showed that older adults reported decreased slow negative responses associated with auditory processing compared to young participants.

Taken together, the largest part of research on the neurophysiology of aging has concentrated on resting state studies. Still, thorough examinations of the age-related neural changes of memory have been produced, pointing to an overall impaired brain functioning in older populations. However, little is known on the impact of aging on the fast-scale brain dynamics underlying predictive and memory processes of sequences unfolding over time. Similarly, the age-related neural changes underlying predictions in cognitive tasks remain elusive.

To address these intriguing topics, the predictive coding theory (PCT) offers a suitable framework. Indeed, PCT states that the brain is constantly updating internal models to predict information and stimuli from the external world ^22^. In recent years, music has emerged as a privileged tool to investigate PCT and understand how the brain encodes, recognises and predicts temporal sequences ^23, 24^. Along this line, in our previous studies we have combined recently developed musical memory paradigms with state-of-the-art neuroimaging techniques, focusing on the brain dynamics of healthy young participants when they encoded and recognised musical sequences. We discovered that encoding of sounds recruited a large network of functionally connected brain areas, especially in the right hemisphere, such as Heschl’s and superior temporal gyri, frontal operculum, cingulate gyrus, insula, basal ganglia, and hippocampus ^25^. Similarly, long-term recognition of short musical sequences recruited nearly the same brain network. However, in this case, the recruitment was bilateral, and it showed hierarchical dynamics from lower- to higher-order brain areas in different frequency bands (e.g. 0.1-1 Hz and 2-8 Hz) ^26^ and in relation to the recognition of previously memorised or varied musical sequences ^27–30^.

In the current study, we took a new fundamental step by using musical memory paradigms and advanced neuroimaging techniques to investigate the impact of aging on the fast-scale brain dynamics underlying predictive and memory processes for musical sequences unfolding over time.

## Results

### Overview of the experimental design and analysis pipeline

In this study, we investigated the impact of aging on the fast-scale spatiotemporal brain dynamics underlying recognition of previously memorised musical sequences. In brief, during magnetoencephalography (MEG) recordings, two groups of participants (39 older adults [older than 60 years old] and 37 young adults [younger than 25 years old]) listened to the first musical sentence of the Prelude in C minor, BWV 847 by Johann Sebastian Bach and were instructed to memorise it to the best of their ability. As shown in **Figure 1** and **Figure S1**, participants were subsequently presented with five-tone musical excerpts (M) taken from the music they previously memorised and with carefully matched variations. The variations consisted of five-tone musical sequences generated by systematically altering the M sequences after either the first (NT1) or third (NT3) tone. For each musical sequence, participants were requested to assess whether the sequence was taken from the memorised musical piece (M) or whether it was new (N). Additional details on the stimuli are available in the Methods section. Key background information on the two samples of participants is reported in **Table 1**.

**Figure 1.**
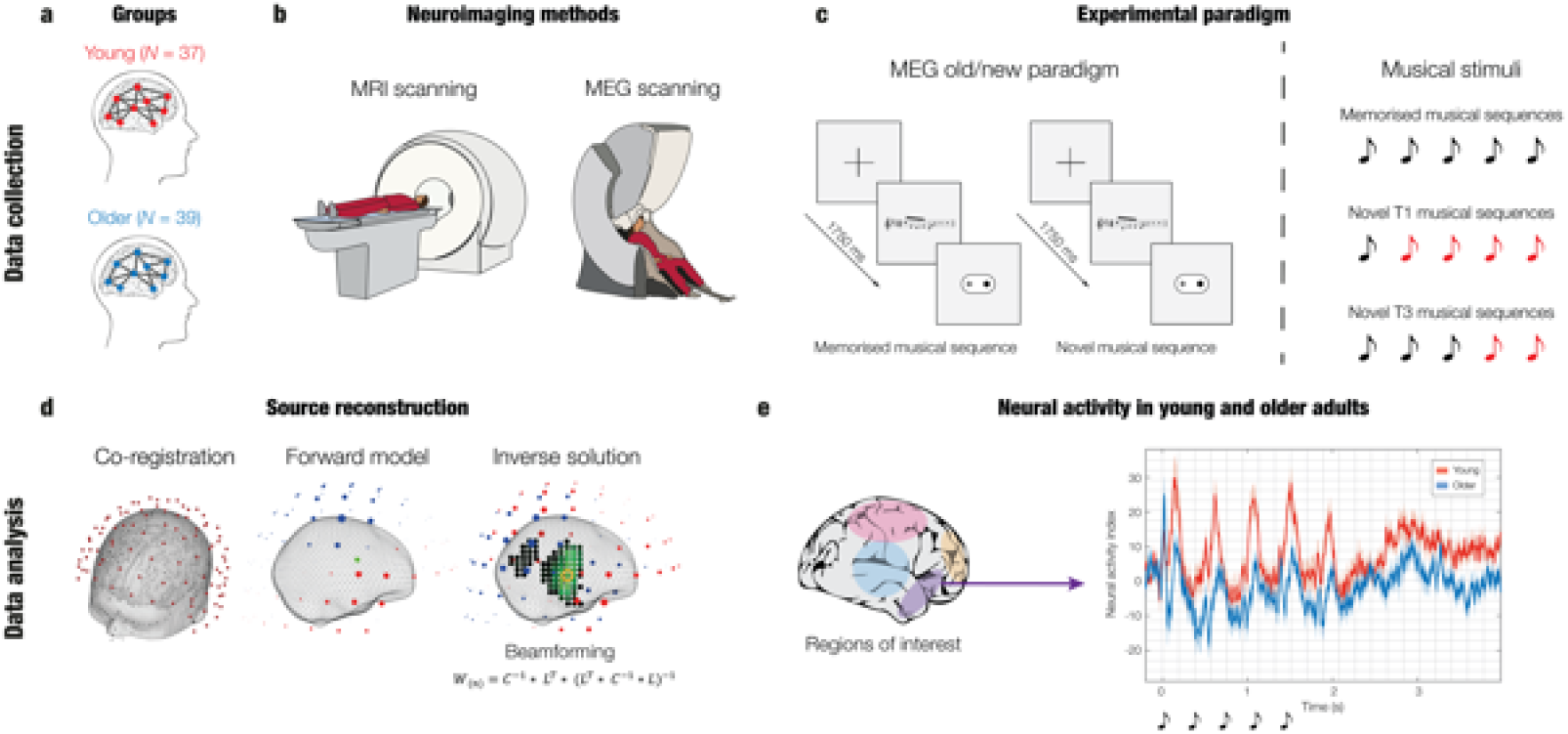
Experimental design, stimuli, and analysis pipeline. **a –** Thirty-seven young and 39 older adults were invited to participate in the experiment. **b –** The brain activity of the participants was collected using magnetoencephalography (MEG), while their structural brain images were acquired using magnetic resonance imaging (MRI). **c –** Participants were requested to memorise a short musical piece (lasting about 30 seconds). Then, we used an old/new auditory recognition task (left). Here, one at a time, five-tone temporal sequences (i.e., musical melodies) were presented in randomised order and participants were instructed to respond with button presses whether they were taken from the musical piece they previously memorised (‘old’ or memorised musical sequences, ‘M’) or they were novel (‘new’ musical sequences, ‘N’). Three types of temporal sequences (M, NT1, NT3) were used in the study. The figure shows a graphical depiction of how the novel musical sequences were created with regards to the previously memorised ones (right). The N sequences were created through systematic variations of the M sequences. For example, in the middle row, it is depicted a sequence (NT1) where we changed all tones but the first one (indicated by the red colours). Likewise, the bottom row shows a sequence where we changed only the last two tones (NT3). **c –** After pre-processing the MEG data, we co-registered it with the individual anatomical MRI data and reconstructed its brain sources using a beamforming algorithm. This procedure returned one time series for each of the 3559 reconstructed brain sources. **e –** We constrained the source reconstructed data to eight brain regions of interest (ROIs) which were selected based on previous literature (left). For each of the ROI, we studied the differences over time between the brain activity of young versus older adults (right).

**Figure 2.**
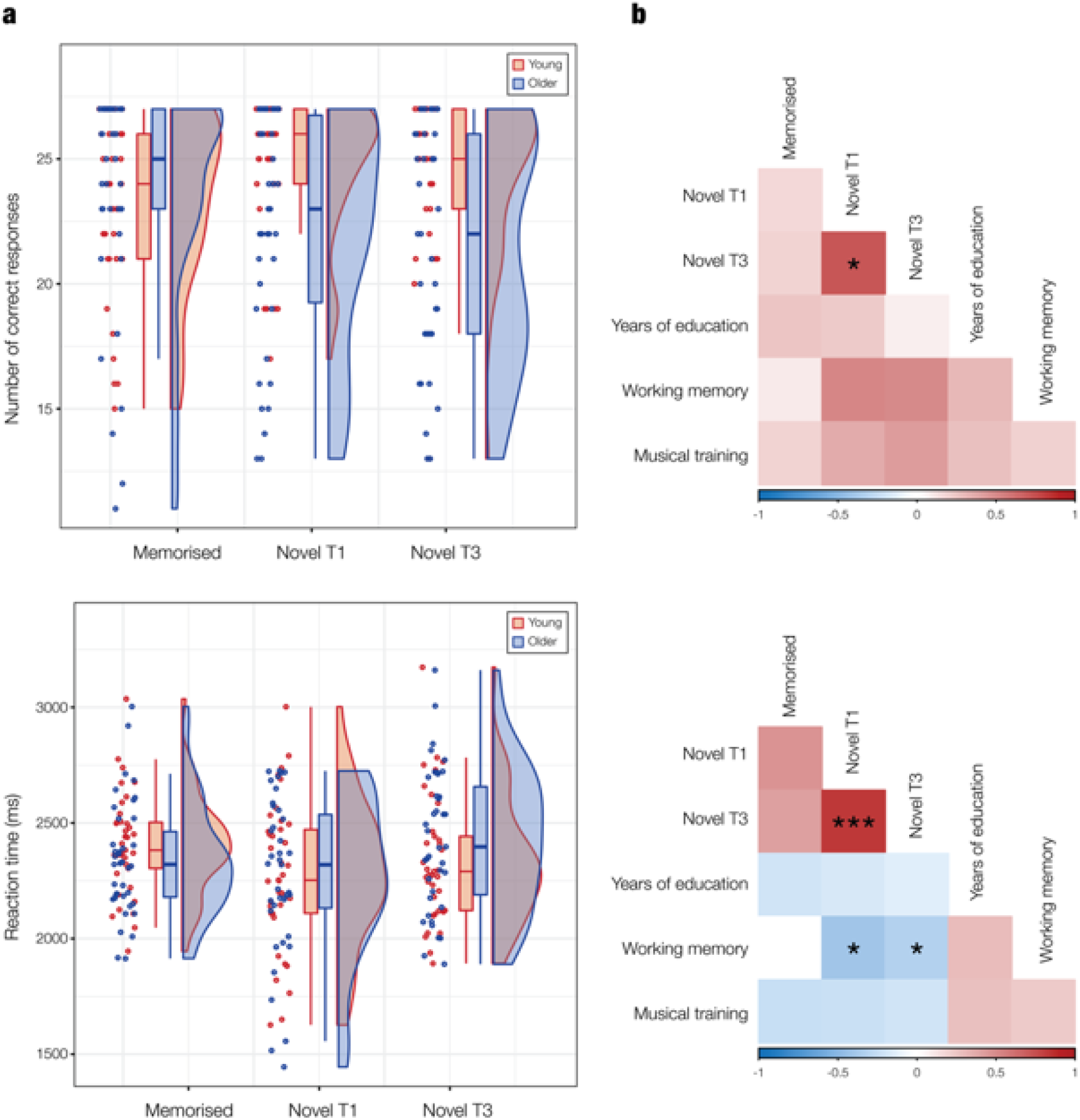
Impact of aging, education, musical training and WM on the recognition of musical sequences. **a –** Raincloud plots show the overlapping distributions and normalized data points of both age groups with regards to the recognition of the previously memorised and novel (NT1 and NT3) musical sequences. Boxplots show the median and interquartile (IQR, 25 – 75%) range, whiskers depict the 1.5*IQR from the quartile. Each dot corresponds to the number of correct responses (top plot) or the mean reaction time (bottom plot) of each participant. The plot above refers to the number accuracy in the task, while the bottom plot to the reaction times. **b –** Correlation matrix between memorised, NT1, NT3 (number of correct responses, top plot, and reaction times, bottom plot), years of education, WM, years of musical training. Significant correlations are indicated by the stars (* p < .05; ** p < .01; *** p < .001).

**Table 1.** Background information on the two age samples. Number of participants, age, sex, WM, years of musical training and general education reported independently for the two age groups. The numbers for age, WM, and years of musical training and general education correspond to mean and standard deviations.

| Participant groups | N | Age | Sex | WM (raw) | Musical training (years) | Education (years) |
| --- | --- | --- | --- | --- | --- | --- |
| Young adults (< 25) | 37 | 21.89 ± 2.05 | 18 F; 19 M | 43.00 ± 7.15 | 3.24 ± 3.72 | 13.57 ± 2.62 |
| Older adults (> 60) | 39 | 67.72 ± 5.35 | 24 F; 15 M | 38.42 ± 7.71 | 3.08 ± 4.31 | 14.20 ± 4.63 |

The analysis pipeline of this study is partly depicted in **Figure 1** and consisted of contrasting the brain activity of young versus older adults at MEG sensor and source levels.

First, we used Monte Carlo simulations (MCS) on univariate tests of MEG sensor data. This was followed by estimating the sources of the brain activity which generated the differences between young and older adults. Second, we focused on eight key regions of interest (ROIs) and analysed whether their time series differed between older and young adults. Third, we assessed the impact of WM, years of general and musical education, sex, and age groups on the brain activity underlying recognition of the musical sequences.

Additional details are available in the Methods section, while the codes used for these analyses are extensively reported at the following links: https://github.com/leonardob92/MEG_Aging_Bach.git

https://github.com/leonardob92/LBPD-1.0.git

### Behavioural results

We calculated the impact of age on response accuracy and reaction times during the musical recognition task that participants performed in the MEG.

Regarding the response accuracy, there was a statistically significant difference between the two age groups in the memory task (*F*(3, 61) = 7.18, *p* < .001, Wilks’ Λ = .739, partial η^2^ = .26). Follow-up ANCOVAs showed that older adults scored lower than young adults when correctly identifying NT1 (*F*(1, 63) = 13.03, *p* < .001) and NT3 sequences (*F*(1, 63) = 19.89, *p* < .001). Years of education (*F*(3, 61) = 3.37, *p* = .02, Wilks’ Λ = .857, partial η^2^ = .14), WM scores (*F*(3, 61) = 7.07, *p* < .001, Wilks’ Λ = .742, partial η^2^ = .26), and years of musical training (*F*(3, 61) = 4.61, *p* = .005, Wilks’ Λ = .815, partial η^2^ = .18) were statistically significant covariates. Specifically, years of education had a statistically significant effect on correctly identifying M (*F*(1, 63) = 4.58, *p* = .03) and NT1 sequences (*F*(1, 63) = 6.52, *p* = .01), meaning that higher number of years of education was associated to higher number of correct responses. Similarly, WM capacity had a statistically significant positive effect on correctly identifying NT1 (*F*(1, 63) = 14.31, *p* < .001) and NT3 sequences (*F*(1, 63) = 19.24, *p* < .001). Finally, years of musical training had a statistically significant positive effect on correctly identifying NT1 (*F*(1, 63) = 5.45, *p* = .02) and NT3 sequences (*F*(1, 63) = 13.80, *p* < .001).

With respect to the average reaction time during recognition of M, NT1 and NT3 sequences, we found a statistically significant difference between the two age groups on the reaction times (*F*(3, 64) = 2.904, *p* = .04, Wilks’ Λ = .880, partial η^2^ = .12). However, this effect was non-significant in follow-up ANCOVAs. Regarding the covariates, only WM scores had a significant effect on the dependent variables (*F*(3, 64) = 5.18, *p* = .002, Wilks’ Λ = .804, partial η^2^ = .20). In particular, we observed that high WM scores were associated with lower average reaction time when correctly identifying NT1 (*F*(1, 66) = 10.96, *p* = .001) and NT3 sequences (*F*(1, 66) = 4.29, *p* = .04).

### Aging and whole-brain activity

To assess the difference between the brain activity of older and young adults while they recognised the musical sequences, we calculated several independent samples t-tests with unequal variances and then corrected for multiple comparisons using cluster-based MCS (t-test threshold = .05, MCS threshold = .001, 1000 permutations). As reported in detail in the Methods section, this procedure was computed independently for the three experimental conditions (M, NT1, NT3).

The analyses returned several significant clusters, highlighting overall reduced brain activity along a wide array of MEG sensors in older participants. In addition, a few significant clusters showed stronger brain activity in older participants. **Table 2** shows the key information of the larger significant clusters for the three experimental conditions, while **Table S1** provides complete statistical information.

**Table 2.**
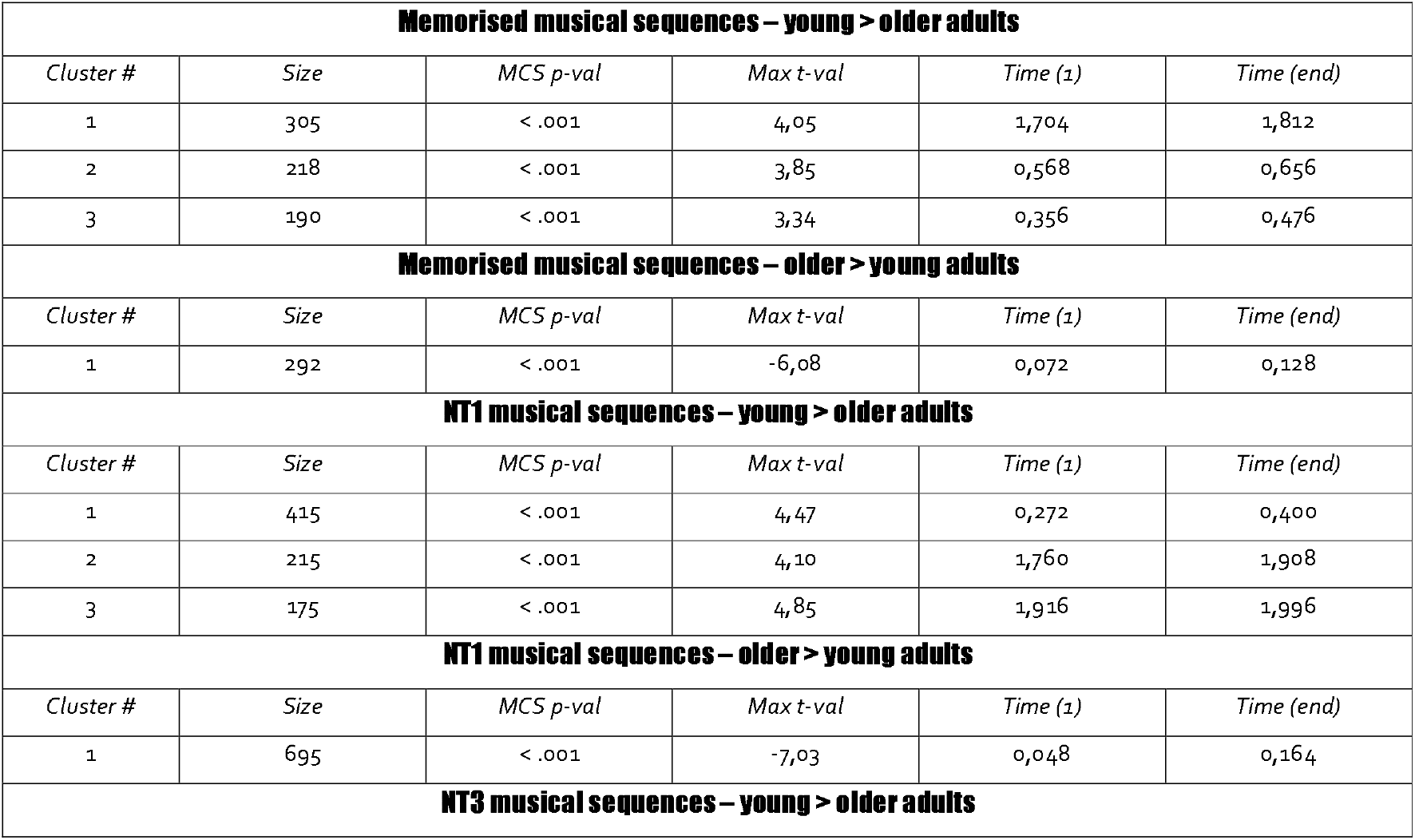
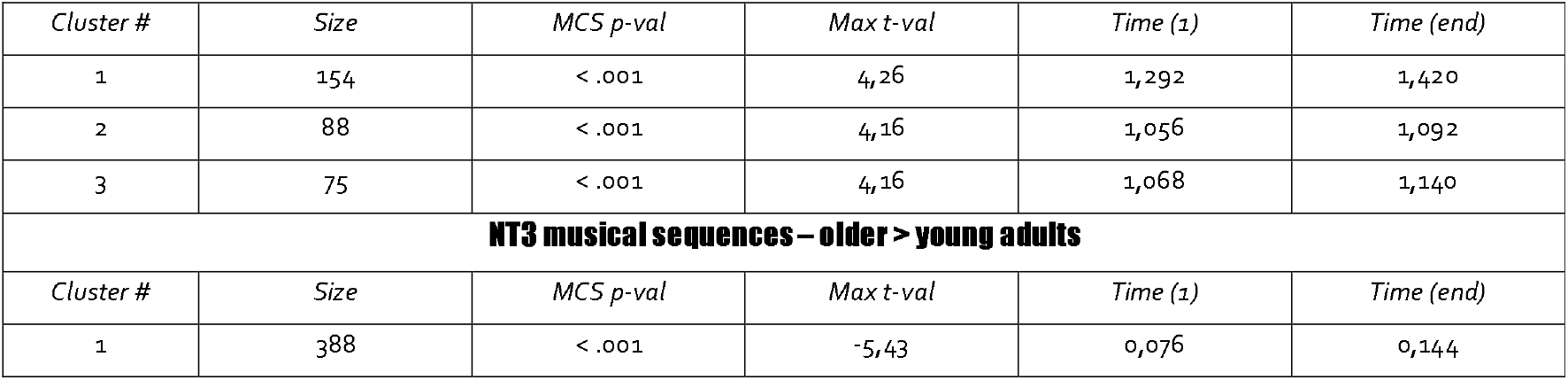
The effect of aging on the whole-brain activity – MEG sensors. Information on the significant clusters emerged at MEG sensors level by contrasting the brain activity underlying recognition of musical sequences of young versus older adults. The results are reported independently for each condition (M, NT1, and NT3) and strength of the contrast (young > older and older > young). The table shows the size of the cluster, the MCS p-value, the maximum t-value within the cluster and the time extent of the significance of the difference between older and young adults.

After analysing the brain activity at the MEG sensor level, we computed source reconstruction analyses using a beamforming algorithm to estimate the brain sources that generated the signal recorded by the MEG sensors. For each of the significant clusters, we contrasted the source-reconstructed brain activity of older versus young adults and corrected for multiple comparisons using a three-dimensional (3D) cluster-based MCS (α < .05, MCS *p*-value = .001). These analyses returned several significant clusters of brain activity, revealing that the main brain regions differentiating older from young adults were the primary and secondary auditory cortices, post-central gyrus, hippocampal regions, inferior frontal gyrus, and ventromedial prefrontal cortex. These results are depicted in **Figures S3** and **S4** and reported in detail in **Table S2** and **S3**.

**Figure 3.**
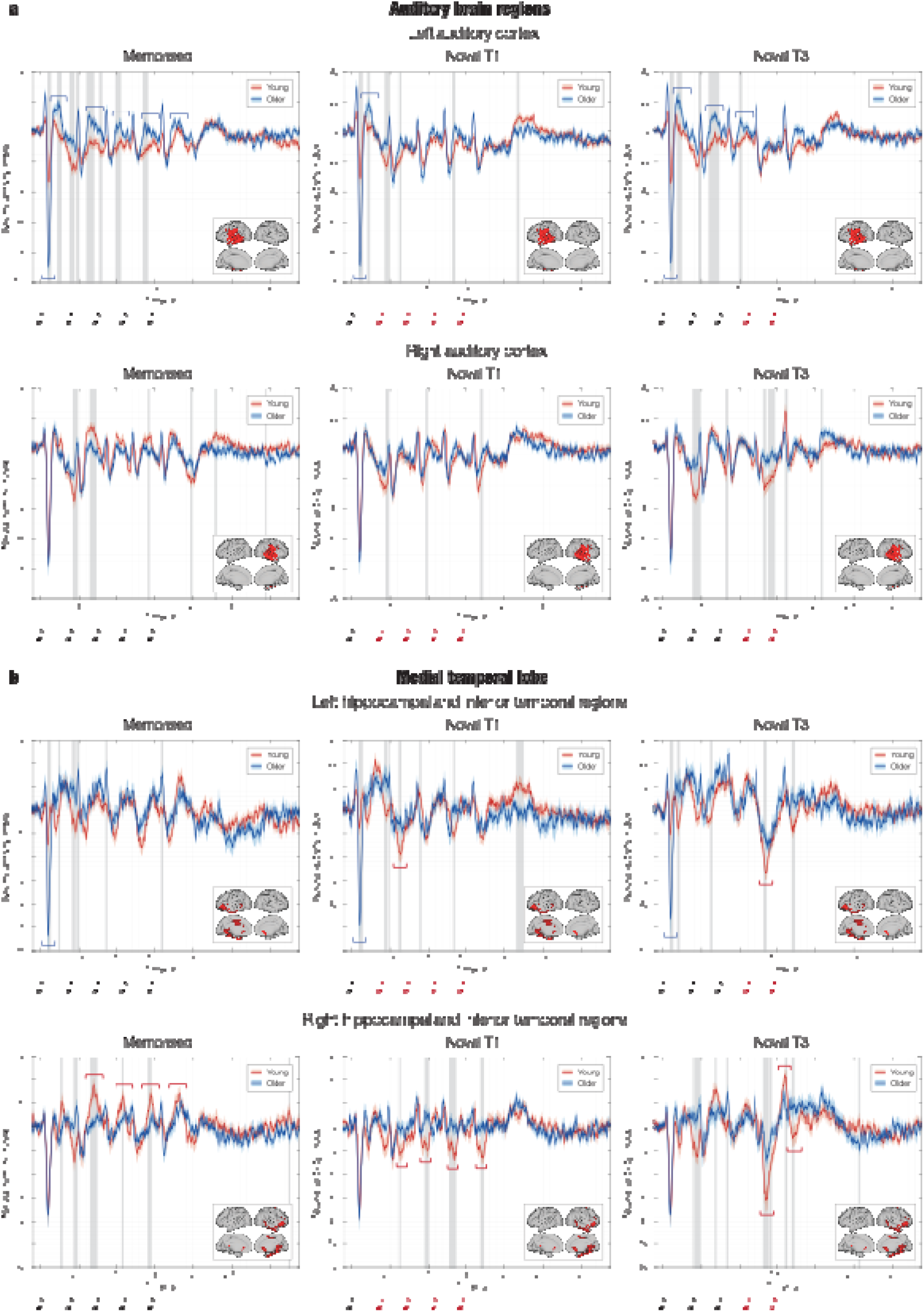
Older adults show stronger activity in auditory cortex and reduced responses in medial temporal lobe during recognition of musical sequences. **a** - The results show that the older adults have significantly stronger activity in the left auditory cortex compared to young adults only when recognising the melodies that were previously memorised. In fact, the top graphs indicate a component occurring about 300 ms after the onset of each tone that was stronger for the older adults for all the tones in the M condition and for all the tones before introducing the variations in the N conditions (i.e. one tone for NT1 and three tones for NT3). In addition, the N100 response to the first tone of the sequences was significantly stronger for old versus young adults in all conditions. **b** - Conversely, older adults showed significantly decreased activity in the hippocampal and inferior temporal regions. This was particularly evident for conditions NT1 and NT3. Here, as highlighted by the red bottom graphs, the older versus young adults exhibited reduced prediction error responses when the sequence was varied. This happened especially for the first tone which introduced the variation in the melodies (i.e. tone two for NT1 and tone four for NT3). Finally, even if to a smaller extent, reduced activity in older adults was also observed for the M condition, where positive components of the neural signals were reduced for all the tones except for the first one. Note that the figure shows the source localised brain activity illustrated for each experimental condition (M, NT1, NT3) in four ROIs (left and right auditory cortex, left and right hippocampal and inferior temporal regions). Grey areas show the statistically significant differences of the brain activity between young (solid red line) and older adults (solid blue, shading indicates standard error in both cases), while red and blue graphs highlight neural components of particular interest. The sketch of the musical tones represents the onset of the sounds forming the musical sequences. The brain templates illustrate the spatial extent of the ROIs.

**Figure 4.**
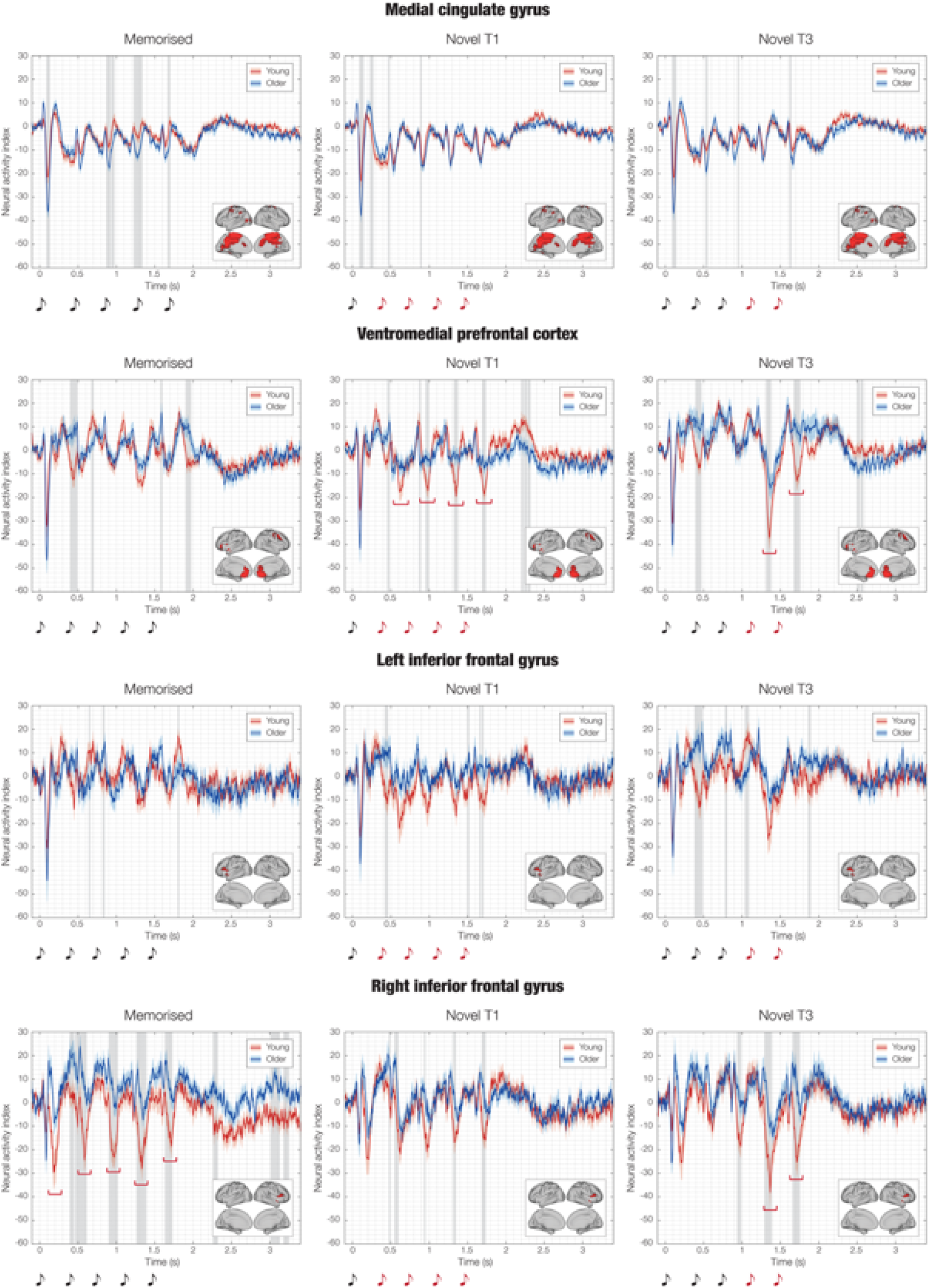
Impact of aging on the cingulate gyrus, ventromedial prefrontal cortex and inferior frontal gyrus responses during recognition of musical sequences. The red graphs in the second row highlight that the VMPFC produced a weaker activity indexing prediction error for the older versus young adults for conditions NT1 and NT3, in an analogous manner to the right hippocampal and inferior temporal regions shown in **Figure 3**. Notably, while these two brain regions also showed a decreased activity for the M condition for older versus young adults, this did not happen for the VMPFC. Finally, the last row of this figure shows a much stronger activity originated in the right inferior frontal gyrus of the young versus older adults. This was particularly evident for the M sequences and consisted of a negative component peaking approximately 250 ms after the onset of each musical tone. Note that the figure shows the source localised brain activity illustrated for each experimental condition (M, NT1, NT3) in four ROIs (medial cingulate gyrus, ventromedial prefrontal cortex [VMPFC], left and right inferior frontal gyrus). Grey areas show the statistically significant differences of the brain activity between young (solid red line) and older adults (solid blue, shading indicates standard error in both cases), while red and blue graphs highlight neural components of particular interest. The sketch of the musical tones represents the onset of the sounds forming the musical sequences. The brain templates illustrate the spatial extent of the ROIs.

### Aging and functional brain regions of interest (ROIs)

To strengthen the reliability of our results and allow an easier comparison with previous literature, we computed a complementary analysis by investigating the difference between the brain activity of older versus young adults in a selected array of functional ROIs that were previously described by Bonetti and colleagues ^28^. These areas (described in detail in **Table S4** and shown in **Figure S2**) were the bilateral medial cingulate gyrus (MC), bilateral ventromedial prefrontal cortex (VMPFC), left (HITL) and right hippocampal area and inferior temporal cortex (HITR), left (ACL) and right auditory cortex (ACR), and left (IFGL) and right inferior frontal gyrus (IFGR). We contrasted the brain activity of young versus older adults by computing an independent-sample t-test for each ROI, timepoint, and condition. We corrected for multiple comparisons using 1D cluster-based MCS (t-value threshold = .05, MCS *p*-value = .001).

This analysis returned several significant clusters showing differences in the brain activity of older compared to young adults. Of particular interest are the clusters reported in the HITR (*p* < .001, k = 25; max *t-val* = 4.70, time: 640 – 736 ms) and IFGR (cluster 1: *p* < .001, k = 38; max *t-val* = −4.59, time: 464 – 612 ms; cluster 2: *p* < .001, k = 33; max *t-val* = −5.04, time: 1260 – 1388 ms) showing reduced activity for older versus young adults when recognising previously memorised musical sequences. In addition, older versus young participants were characterised by a weaker signal in response to the variation of the original musical sequences. This was particularly evident for HITR (NT1: *p* < .001, k = 24; max *t-val* = −3.53, time: 1284 – 1376 ms; NT3: *p* < .001, k = 21; max *t-val* = −4.01, time: 1320 – 1400 ms), VMPFC (NT1: *p* < .001, k = 15; max *t-val* = −3.57, time: 1320 – 1376 ms; NT3: *p* < .001, k = 23; max *t-val* = −3.97, time: 1672 – 1760 ms), and HITL (NT3: *p* < .001, k = 12; max *t-val* = - 3.31, time: 1324 – 1368 ms).

Finally, older adults showed a stronger activity in ACL in response to the first tone of the sequences in all conditions (M: *p* < .001, k = 14; max *t-val* = 5.37, time: 84 – 136 ms; NT1: *p* < .001, k = 15; max *t-val* = 5.86, time: 88 – 144 ms; NT3: *p* < .001, k = 16; max *t-val* = 6.04, time: 84 – 144 ms) and in relation to each tone until the variation was introduced (**Figure 3**, first row). These results are depicted in **Figures 3** and **4** and extensively reported in **Table S5**.

### WM, musical expertise, education level, aging and neural data

Finally, we computed two additional analyses to assess whether potential confounding variables had an impact on the relationship between aging and the neural mechanisms underlying recognition of musical sequences.

In the first analysis we computed three independent multivariate analyses of covariance (MANCOVAs), one for each experimental condition. In each MANCOVA, the dependent variables were the highest peaks of the neural data for the eight ROIs, while the independent variables were age, sex, years of formal musical expertise, WM, and years of formal education (see Methods for additional details).

The results of the MANCOVAs showed a significant main effect for age in all experimental conditions: M (*F*(8, 59) = 4.62, *p* = .0002, Wilks’ Λ = .614, partial η^2^ = .39), NT1 (*F*(8, 59) = 3.117, *p* = .005, Wilks’ Λ = .703, partial η^2^ = .30), and NT3 (*F*(8, 59) = 3.575, *p* = .002, Wilks’ Λ = .674, partial η^2^ = .33). This confirmed the impact of age on the neural data. The other variables did not show any significant results, indicating that no confounding variables affected the relationship between age and the neural data. However, WM approached the significance in all experimental conditions, showing moderate effect sizes: M (*F*(8, 59) = 4.62, *p* = .09, Wilks’ Λ = .802, partial η^2^ = .20), NT1 (*F*(8, 59) = 1.313, *p* = .25, Wilks’ Λ = .849, partial η^2^ = .15), and NT3 (*F*(8, 59) = 1.691, *p* = .11, Wilks’ Λ = .814, partial η^2^ = .19). This indicated that WM may partially affect the brain dynamics of musical recognition in relation to aging.

Following the results of the MANCOVAs, we computed independent analyses of variance (ANOVAs) for each time-point, ROI and condition and used cluster-based 3D MCS to correct for multiple comparisons. We used two-way ANOVAs with the following levels: WM (high and low performers) and age (older and young adults). The analysis returned significant key clusters for three main ROIs in the NT3 condition: HITR (NT3: *p* < .001, k = 40; max *F-val* = 17.66, time: 1308 - 1464 ms), VMPFC (NT3: *p* < .001, k = 32; max *F-val* = 13.57, time: 1300 - 1424 ms), HITL (NT3: *p* < .001, k = 24; max *F-val* = 9.36, time: 1304 - 1396 ms). **Figure 5** show the time series of these ROIs in relation to age and WM, while detailed statistical results are reported in **Table S6**.

**Figure 5.**
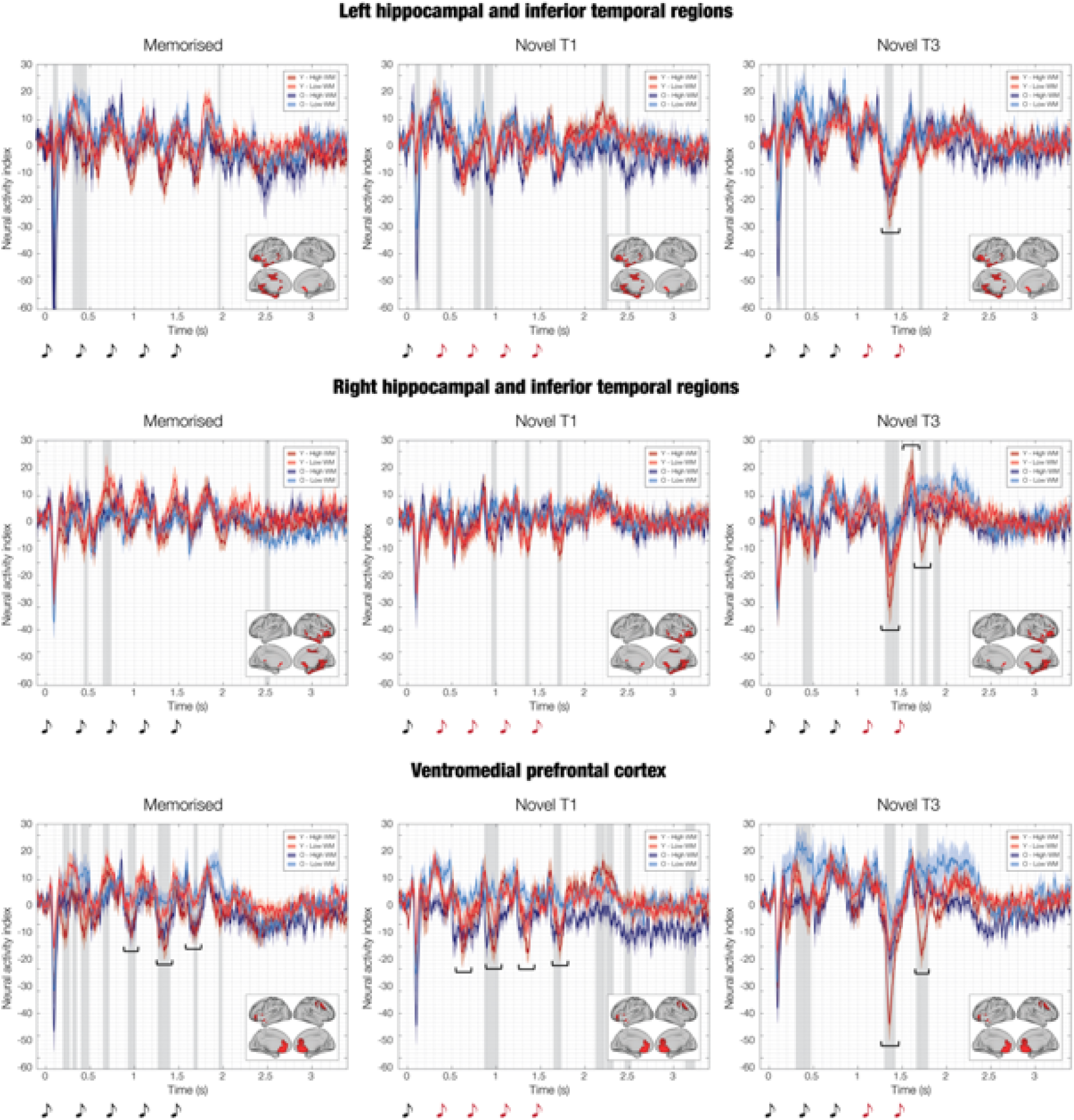
Impact of WM and aging on the ventromedial prefrontal cortex and medial temporal lobe responses during recognition of musical sequences. The black graphs in the NT3 plots (all rows) highlight that the strongest brain prediction error in response to the variation of the original musical sequences occurred in young adults who performed very well in the WM tasks. The strength of the prediction error was lower and very similar for young adults with low WM and older adults with high WM. Finally, older adults with low WM presented the most reduced prediction error signal in the brain. This was particularly evident for the right hippocampal and inferior temporal regions as well as for the VMPFC. A similar, but less pronounced, effect was observed in the VMPFC for M and NT1. Note that the figure shows the source localised brain activity illustrated for each experimental condition (M, NT1, NT3) in three (ventromedial prefrontal cortex [VMPFC], left and right hippocampal and inferior temporal regions). Graphs indicates the key event of interest in the brain responses, while the grey areas show the statistically significant differences of the brain activity between the participants grouped in the following four groups: young adults - high WM (i), young adults - low WM, older adults - high WM, older adults - low WM. Solid line indicates the average over participants, independently for the four groups, while the shaded area the standard errors. The sketch of the musical tones represents the onset of the sounds forming the musical sequences. The brain templates illustrate the spatial extent of the ROIs.

Finally, we computed an additional sub-analysis to assess whether we could distinguish a sub-sample of the older participants based on their brain activity. To this aim, we used one-way ANOVAs contrasting three age-groups: young (younger than 25), older adults 60-68 (age between 60 and 68, n = 23) and older adults > 68 (older than 68, n = 16). Then, we corrected for multiple comparisons with cluster-based 3D MCS. The results highlighted that the oldest group within the older adults was characterised by overall reduced brain activity, especially in response to the variation of the original sequences. This was particularly evident for HITR (NT3: *p* < .001, k = 22; max *F-val* = 7.92, time: 1312 - 1396 ms), VMPFC (NT3: *p* < .001, k = 15; max *F-val* = 7.73, time: 1320 - 1376 ms), HITL (NT3: *p* < .001, k = 13; max *F-val* = 10.01, time: 1316 - 1364 ms). **Figure 6** shows the time series of these ROIs in relation to the three age groups, while detailed statistical results are reported in **Table S7**.

**Figure 6.**
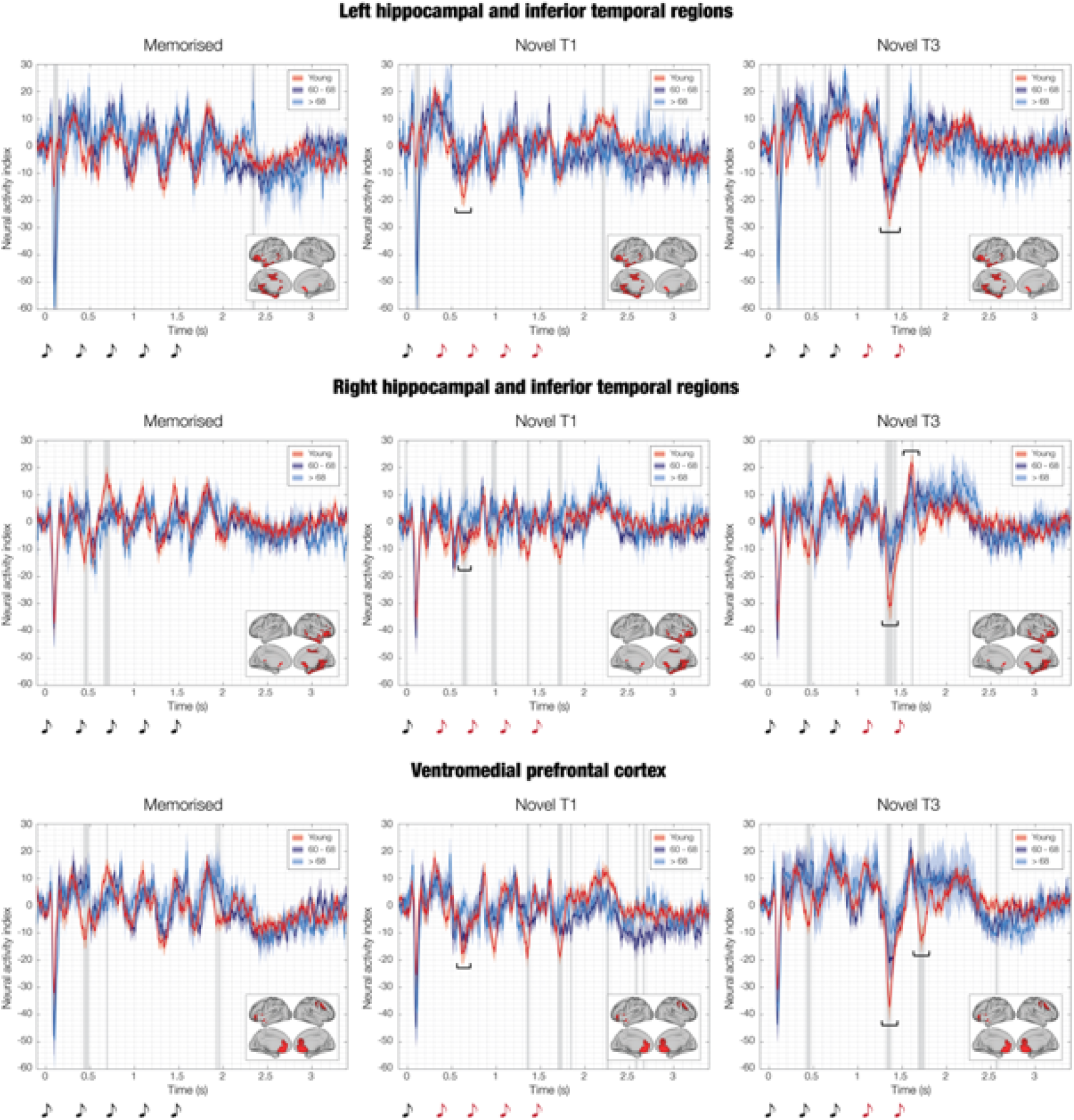
Ventromedial prefrontal cortex and medial temporal lobe responses during recognition of musical sequences for three age groups (young adults, adults between 60 and 68 years of age, adults older than 68). The black graphs in the NT1 and NT3 plots (all rows) highlight that the strength of the brain prediction error in response to the variation of the original musical sequences was modulated by age. In fact, the strongest signal was recorded for the young adults. A reduced prediction error was observed for the adults aged between 60 and 68, while the weakest signal occurred for the adults older than 68 years. As observed for the WM in **Figure 5**, this effect was particularly evident for the VMPFC and right hippocampal and inferior temporal regions. Note that the figure shows the source localised brain activity illustrated for each experimental condition (M, NT1, NT3) in three (ventromedial prefrontal cortex [VMPFC], left and right hippocampal and inferior temporal regions). Graphs indicates the key event of interest in the brain responses, while the grey areas show the statistically significant differences of the brain activity between the participants grouped in the following three groups: young adults (i), adults between 60 and 68 years of age (ii), adults older than 68 years (iii). Solid line indicates the average over participants, independently for the four groups, while the shaded area the standard errors. The sketch of the musical tones represents the onset of the sounds forming the musical sequences. The brain templates illustrate the spatial extent of the ROIs.

## Discussion

In this study, we have combined fast neuroimaging with the recognition of previously memorised and novel musical sequences to unpick the complexity of the healthy aging brain. Our findings challenge simplistic notions that non-pathological aging merely diminishes neural predictive capabilities by showing age-related neural transformations in predictive and memory processes.

During the recognition of the previously memorised melodies, the left auditory cortex exhibited stronger activity in response to each sound of the sequence for the older compared to young adults. Conversely, other brain regions of key importance for memory and predictive processes such as the hippocampus, inferior temporal cortex and inferior frontal gyrus showed an overall decreased activity for the older adults. In response to the varied musical sequences, the left auditory cortex did not exhibit any difference between older and young adults after the musical sequence was altered. Conversely, a much-reduced activity was observed for the older adults after the sequence was changed. This effect was particularly strong for the condition NT3 where the sequence was altered after the fourth tone, and it primarily regarded hippocampus, inferior temporal cortex and ventromedial prefrontal cortex. Working memory (WM) abilities also affected the brain responses, especially for the condition NT3, both in older and young individuals. The brain activity after varying the original musical sequence was reduced for participants with lower WM skills.

In relation to the behavioural responses, no differences between older and young adults were found when inspecting the accuracy and reaction times associated with the recognition of the previously memorised sequences. Conversely, older adults reported lower accuracy when recognising the varied musical sequences (both NT1 and NT3). No differences were observed for the reaction times.

As expected, the results of this study are consistent with our previous research on the brain dynamics underlying the encoding and recognition of musical sequences in healthy young individuals, which showed that the recognition of the previously memorised and varied musical sequence is built over time through a rapid hierarchical pathway of components originated in the auditory cortex and progressing to the hippocampus, ventromedial prefrontal cortex and inferior temporal cortex ^25–30^. Beyond this, the most notable finding of our study is the altered brain functioning observed in older compared to young adults. On the one hand, this occurred through an overall reduction of the brain activity generated in memory brain regions, supporting previous findings which reported diminished brain responses in aging populations in a variety of different contexts, spanning from resting state to automatic neural responses and conscious tasks ^5, 6, 8, 13, 14, 17, 20, 31^. On the other hand, only for the recognition of the previously memorised sequences, older adults showed increased activity in the left auditory cortex. This altered brain functioning supports the hypothesis that neural predictive processes in non-pathological aging are not simply reduced, but qualitatively transformed ^32^.

Overall, our results can be very well interpreted within the large framework of the PCT, providing a relevant contribution to the age-related changes of its neural underpinnings. PCT posits that the brain is constantly updating internal models to predict information and stimuli from the external world ^22^. Recently, it has been successfully linked to complex cognitive processes, finding a notable example in the neuroscience of music. Vuust and colleagues ^23, 24^ suggested that, while processing music, the brain repeatedly generates hypotheses and predictions about the upcoming unfolding of musical sequences. When the prediction matches the incoming sounds, the brain recognises the music. Conversely, when the expectation is violated by different sounds, predictions errors arise. Our findings point to impaired conscious predictive coding processes in healthy older adults, as evidenced by reduced brain activity during the prediction and recognition of both original and varied musical sequences. As such, they provide novel insights into the brain dynamics of PCT across the lifespan of healthy adults. These results are also coherent with previous studies which showed an age-related reduction of automatic predictive processes such as MMN ^13–15^. Notably, our study largely expands on their significance by showing age-related changes of conscious predictive processes and novelty detection and not only automatic responses to subtle environmental irregularities as typically done in MMN studies.

Along this line, we revealed decreased activity in older adults during the recognition of the previously memorised musical sequences in brain regions particularly relevant for memory and predictive processes, such as the hippocampus (especially in the right hemisphere) ^33, 34^. Numerous studies have shown the detrimental effects of aging on the hippocampus and memory performance. For instance, it has been shown that aging is associated with reduced hippocampal size ^35, 36^ and that it affects the long-term potentiation (LTP) and long-term depression (LTD) occurring in the hippocampal neurons ^37^. The altered size and functionality of LTP and LTD in the hippocampus occurring with aging might be reflected in the reduction of hippocampal activity that we observed in older adults in our study during the recognition of the previously learned musical sequences. The stronger involvement of the hippocampus in the right hemisphere is coherent with the plethora of findings which reported the right-hemispheric dominance in music processing ^38^.

Another essential brain region for understanding, predicting and producing language and music is the inferior frontal gyrus ^39, 40^, which also showed a sharp decreased activity in older adults in our study. This is a rather interesting result since there is scarce evidence showing impaired functionality in the inferior frontal gyrus in aging populations, suggesting that this effect may be specifically linked to the age-related changes underlying the prediction and recognition of musical sequences. Moreover, the inferior frontal gyrus does not normally play a pivotal role in the recognition of memorised musical sequences. However, our results suggest that it may provide an additional, relevant contribution to this memory process, which is instead largely attenuated in aging populations, as clearly shown by the contrast between the brain activity of young and older adults. This evidence might also point to a general reduced functionality of the inferior frontal gyrus in older adults, potentially contributing to explain the challenges older adults often face in linguistic, predictive and memory tasks ^41–43^.

On a related note, our findings revealed an intriguing pattern of increased activity in the left auditory cortex of older adults during the recognition of musical sequences. This increased activity was observed for the N100 component to the first sound, as well as for the positive component peaking around 350-400 ms after each sound of the sequence. Coherently with PCT, it is plausible that the increased activity in the left auditory cortex is a result of top-down influences from the hippocampus and ventromedial prefrontal and inferior temporal cortices, which are supposed to actively monitor the unfolding musical sequence ^24^. In this case, when they successfully predict the sequence, they require less effort from the left auditory cortex. In a young and more effective brain, the more refined prediction and higher control exerted by those brain regions would result in a reduced activity in the auditory cortex, exactly as we observed in our study.

Interestingly, no significant differences were found between older and young adults in terms of accuracy and reaction times when recognising previously memorised sequences. This finding suggests that brain activity may undergo alterations before behavioural manifestations become apparent. This observation raises exciting possibilities for using brain activity as a potential biomarker for the early detection of cognitive decline, which should be further explored by future studies.

We also examined the impact of aging on the recognition of varied musical sequences and the prediction error arising when the original sequences were altered. We identified the key involvement of the left and, especially, right hippocampus and bilateral ventromedial prefrontal cortex. The hippocampus is a central brain region for prediction error ^34, 44^ and its reduced activity in older adults suggests that aging is associated with decreased ability to consciously process errors and deviations from previously learned sequences.

Similarly, the ventromedial prefrontal cortex, a brain region implicated in reasoning and evaluation processes ^45^, exhibited reduced activity in older adults. In accordance with our findings, studies have shown that age-related changes in the ventromedial prefrontal cortex are associated with decline in cognitive control and decision-making abilities ^46, 47^. For instance, O’Callaghan and colleagues ^46^ found that individuals with ventromedial prefrontal cortex damage and healthy older adults reported reduced awareness of the presented stimuli during learning tasks. This relates to our results, suggesting that the reduced activity in the ventromedial prefrontal cortex observed in older adults might represent the neural signature of the decreased conscious prediction error and awareness of the musical novelty in aging.

To be noted, splitting the older adult participants into two age groups further strengthens the reliability of our previously described results, as it reveals a more pronounced reduction in brain activity in participants older than 68 compared to those aged 60-68. This highlights the progressive nature of age-related changes in brain functioning.

Lastly, we showed a relationship between the participants’ WM abilities and the brain activity. Participants with higher WM exhibited stronger brain activity, particularly when recognising the varied musical sequences. This finding underscores the potential of using WM as a predictor of preserved brain activity in older adults. In fact, older adults with high WM capacity showed brain activity levels similar to those of young adults with lower WM capacity. This finding is strongly in line with previous research on cognitive reserve, suggesting that higher cognitive abilities in older populations represent a protective factor against mild cognitive impairment and dementia ^48–50^.

In summary, the present study provides valuable novel insights into the impact of aging on the brain function and shows how age is not always related to decline but rather to a comprehensive transformation of brain regions, including the hippocampus, inferior frontal gyrus, and ventromedial prefrontal and auditory cortices. The results provide an important contribution to understanding age-related neural changes and reveal the potential of our methods to identify possible biomarkers for healthy aging and early detection of transformative changes in brain function.

## Materials and methods

### Participants

After removing one participant due to technical issues with the MEG signal, the sample consisted of 76 participants (34 males, 42 females), divided into two age groups: young and older adults. The older adult group consisted of 39 participants (24 females, 15 males) aged 60 to 81 years old (mean age: 67.72 ± 5.35 years). The young group included 37 participants (18 females, 19 males) aged 18 to 25 years old (mean age: 21.89 ± 2.05 years). The nationality of all participants was Danish. The inclusion criteria for the participants were the following: (i) normal health (no reported neurological nor psychiatric illness), (ii) age between 18 and 25 years old (young adults’ group) and older than 60 years (older adults’ group), (iii) normal hearing according to the age group of each participant, (iv) normal sight or corrected to normal sight (e.g., contact lenses), and (v) understanding and acceptance of participant information. The exclusion criteria that we applied were: (i) use of prescribed medication that could affect the central nervous system, (ii) neurological or psychiatric illness, (iii) lack of cooperation or verbal agreement for participating in the study, (iv) magnetic resonance imaging (MRI) contraindications, (v) age between 26 and 59 years old, and impaired hearing (vi).

The project was approved by the Institutional Review Board of Aarhus University (case number: DNC-IRB-2021-012). The experimental procedures complied with the Declaration of Helsinki – Ethical Principles for Medical Research. Participants’ informed consent was obtained before the beginning of the experiment.

### Experimental stimuli and design

In this study, we presented participants with an auditory recognition task based on the old/new paradigm that we developed in our previous works ^26–30^. At the same time, we recorded their brain activity using magnetoencephalography (MEG). The participants were required to listen to a brief musical piece (roughly 25 seconds) twice and were instructed to memorise it as best as they could. The musical piece comprised the initial four measures of Johann Sebastian Bach’s Prelude No. 2 in C Minor, BWV 847. The wave audio file that we used in the experiment was generated using Finale (MakeMusic, Boulder, CO). The volume of the musical stimuli was set to 60 dB for 67 participants and to 70 dB on average for nine of our participants older than 70 years who presented a very mild hearing impairment, as typically occurring with aging. To limit the adjustment of the volume across participants to only a few of them, we used sounds that almost always fell in the range 125 - 650 Hz, which is only marginally affected by the typical hearing loss occurring with aging ^51^. Each tone within the piece had the same duration of around 350 ms. In the second phase of the task, participants were presented with 81 musical sequences consisting of five tones and lasting 1750 ms. They were then asked to identify whether each sequence was part of the original musical piece (old or memorised sequence [M]) or if it was a different musical sequence (new or novel sequence [N]) (see **Figure 1**). For the purpose of this study, we presented participants with 27 sequences from the original musical piece and created 54 variations of the original melodies. The musical sequences used in the study are depicted in **Figure S1**. The two types of stimuli used in the study were created as follows. The M sequences were comprised of the first five tones from the first three measures of the musical piece. These sequences were presented a total of 27 times, nine times for each sequence. The N sequences were generated by systematically altering the three M sequences (see **Figure 1**). This involved changing every musical tone of the sequence while keeping the first tone (NT1) or the first three tones (NT3) the same as the M sequences. Nine variations were created for each of the original M sequences and each of the two categories of N. As a result, there were 27 N sequences for each category and 54 N sequences in total. The variations were created following specific rules:

- Inverted melodic contour (used twice): this involved creating a variation with a melodic contour that was inverted relative to the original M sequence. (i.e., if the melodic contour of the M sequence was down-down-up-down, the N sequence would be up-up-down-up).
- Same tone scrambled (used three times): this involved scrambling the remaining tones of the M sequence (e.g., M sequence C-E-D-E-C, was changed into NT1 sequence C-C-E-E-D).
- Same tone (used three times): this involved using the same tone repeatedly, sometimes varying only the octave (e.g., M sequence C-E-D-E-C, became NT1 sequence C-E^8-^ E^8-^ E_8_^-^ E_8_).
- Scrambling intervals (used once): this involved scrambling the intervals between the tones (e.g., M sequence 6^th^m - 2^nd^m – 2^nd^m – 3^rd^m, was changed to NT1 sequence 2^nd^m, 6^th^m, 3^rd^m, 2^nd^m).

We adopted this procedure to study the difference between young and older adults with regards to their brain dynamics underlying *(i)* the recognition of previously memorised auditory sequences and (*ii*) their conscious detection of the varied sequences.

### Neural data acquisition

During this study, MEG recordings were conducted at Aarhus University Hospital (AUH), Aarhus, Denmark, using an Elekta Neuromag TRIUX MEG scanner with 306 channels. The data was recorded with an analogue filtering of 0.1 – 330 Hz at a sampling rate of 1000 Hz. To ensure accurate co-registration with the MRI anatomical scans, the head shape of participants and the position of four Head Position Indicator (HPI) coils were registered using a 3D digitizer (Polhemus Fastrak, Colchester, VT, USA). During the MEG recordings, two sets of bipolar electrodes were also used to record cardiac rhythm and eye movements, allowing for removal of electrocardiography (ECG) and electro-oculography (EOG) artifacts in a later stage of the analysis.

The MRI scans were recorded on a CE-approved 3T Siemens MRI-scanner at AUH using the following structural T1 sequence parameters: echo time (TE) = 2.61 ms, repetition time (TR) = 2300 ms, reconstructed matrix size = 256 x 256, echo spacing = 7.6 ms, and bandwidth = 290 Hz/Px.

The MEG and MRI recordings were conducted on separate days.

### Working memory, musical expertise and background data

We evaluated domain-general working memory (WM) abilities using the Digit Span and Arithmetic subtests from the Wechsler Adult Intelligence Scale IV’s Working Memory index. The Digit Span subtest required participants to listen and repeat sequences of numbers in the same, inverse, or ascending order. The Arithmetic subtest involved solving mathematical operations provided orally by the experimenters without external aids. We combined the raw scores from both subtests to calculate individual WM abilities, with scores ranging from five to 70. Additionally, we assessed formal musical training using the Goldsmiths Musical Sophistication Index (Gold-MSI) questionnaire, which includes 39 questions on musical skills, experience, and habits. We used the Musical Training facet, which estimates an individual’s history of formal musical training, and scores range from seven to 49.

In addition, we collected general background data such as the years of education. These data were then used as covariates in later stages of the analysis to assess whether they had an impact on the relationship between age and neural data during recognition of auditory sequences.

### Behavioural data during MEG recording

During the auditory recognition task, we recorded participants’ responses and reaction times. We then used this data to estimate differences in response accuracy and average reaction time between young and older participants, and to calculate the impact of sex, years of education, WM abilities, and years of musical training on the behavioural data.

We computed two independent multivariate analysis of variance (MANCOVA, Wilk’s Lambda [Λ], α = .05) ^52^ using group as the independent variable (young vs older) and years of education, WM scores, years of musical training, and sex as covariates. In one MANCOVA, number of correct responses (divided into M, NT1 and NT3) were used as the three dependent variables. In the other MANCOVA, average reaction time during correct responses (divided into M, NT1, and NT3) were used as the three dependent variables. The effect size was calculated using partial eta squared (i.e., partial η^2^).

To determine the effects of the independent variable and covariate, univariate analyses of covariance (ANCOVA) were computed individually for each of the dependent variables and statistically significant covariates.

### MEG data pre-processing

The MEG data obtained from 204 planar gradiometers and 102 magnetometers was initially subjected to pre-processing with MaxFilter ^53^, which helped to reduce external interferences. We applied signal space separation and the following MaxFilter parameters: spatiotemporal signal space separation [SSS], down-sample from 1000Hz to 250Hz, correlation limit between inner and outer subspaces used to reject overlapping intersecting inner/outer signals during spatiotemporal SSS: 0.98, movement compensation using cHPI coils (default step size: 10 ms).

After conversion to Statistical Parametric Mapping (SPM) format, the data was pre-processed and analysed in MATLAB using both in-house-built codes (LBPD, https://github.com/leonardob92/LBPD-1.0.git) and the freely available Oxford Centre for Human Brain Activity (OHBA) Software Library (OSL) ^54^ (https://ohba-analysis.github.io/osl-docs/), which utilises Fieldtrip ^55^, FSL ^56^, and SPM ^57^ toolboxes. We visually inspected the filtered MEG data using OSLview to remove large artifacts, which accounted for less than 0.1% of the total data. We employed independent component analysis (ICA) to separate and remove eyeblink and heartbeat interference from the brain data ^58^. This involved decomposing the original signal into independent components, discarding the components that detected eyeblink and heartbeat activities, and reconstructing the signal using the remaining components. We then epoched the signal into 81 trials and baseline-corrected it by subtracting the mean signal recorded in the baseline from the post-stimulus brain signal. The trials lasted 3500 ms (3400 ms after the onset of the first tone of the musical sequence plus 100 ms of baseline time) and were categorised into three groups (M, NT1, NT3) with 27 trials each.

### MEG sensor level and aging

To assess the difference between the brain activity of young and older adults while they recognised the musical sequences, we calculated several independent samples t-tests with unequal variances and then corrected for multiple comparisons using cluster-based Monte-Carlo simulations (MCS). As it is common in MEG and EEG task studies ^59, 60^, we computed the average over trials in each condition before performing t-tests, which resulted in three mean trials (M, NT1, NT3). For each condition separately, we computed a t-test for each MEG magnetometer channel and each time-point between 0 and 2000 ms, contrasting the brain activity of young and older adults. We then reshaped the matrix to obtain a two-dimensional (2D) approximation of the MEG channels layout for each time-point, binarising it based on the *p*-values obtained from the previous t-tests (threshold = .05) and the sign of t-values. The resulting 3D matrix (*MX*, 2D x time) consisted of 0s when the t-test was not significant and 1s when it was. To correct for multiple comparisons, we identified clusters of 1s and assessed their significance using MCS. Specifically, we performed 1000 permutations of the elements of the original binary matrix *MX*, identified the maximum cluster size of 1s, and built the distribution of the 1000 maximum cluster sizes. We considered clusters that had a size bigger than the 99.9% maximum cluster sizes of the permuted data to be significant. We applied the MCS procedure to the absolute values of magnetometer MEG channels for both young versus older adults and vice versa.

### Source reconstruction

MEG provides excellent temporal resolution, but to fully understand the brain activity underlying complex cognitive tasks, the spatial component of the brain activity must also be identified. To estimate the sources of the brain that generated the signal recorded by the MEG sensors, we computed a source reconstruction protocol using a combination of in-house-built codes and codes available in OSL, SPM, and FieldTrip.

The source reconstruction analysis consists of designing a forward model and computing the inverse solution. The forward model considers each brain source as an active dipole and describes how the unitary strength of each dipole is reflected over all MEG sensors. We used magnetometer channels and an 8-mm grid to obtain 3559 dipole locations within the whole brain (voxels). After co-registering the individual structural T1 data with the fiducial points (i.e., information about head landmarks such as the nasion and the left and right pre-auricular points), we computed the forward model using the widely used “Single Shell” method, which resulted in a leadfield model stored in matrix *L* (sources x MEG channels) ^61^. In cases where structural T1 was unavailable, we used a template (MNI152-T1 with 8-mm spatial resolution) for the leadfield computation.

Afterwards, we calculated the inverse solution, using the established beamforming method, which is a popular and effective algorithm available in the field of neuroscience. The process involves utilising a distinct series of weights that are applied successively to the source positions, enabling the separation of the impact of each source on the activity detected by the MEG channels. This is carried out for every instance of the brain data captured. The beamforming inverse solution is comprised of several key stages, which can be outlined as follows.

The data measured by the MEG sensors (*B*) at time *t*, can be described by the following equation (1):

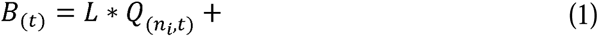

where *L* is the leadfield model, *Q* is the dipole matrix which carries the activity of each active dipole (*q*) over time, and LJ is noise (see Huang and colleagues for details ^62^). In order to resolve the inverse problem, *Q* has to be computed. In the beamforming algorithm, to calculated *Q*, a series of weights have to be computed and applied to the MEG sensors at each timepoint. This is done for each single dipole *q* and shown in equation (2):

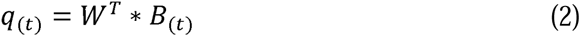

To obtain *q*, the weights *W* have to be computed (here, the subscript *T* indicates the transpose matrix). The beamforming method relies on the matrix multiplication between *L* and the covariance matrix between MEG sensors (*C*). This matrix is calculated on the concatenated experimental trials. More specifically, for each brain source *n*, the weights *W_n_* are calculated as shown in equation (3):

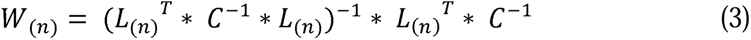

The calculation of the leadfield model was performed for the three main orientations of each brain source (dipole), as done in the field (see, for example, Nolte ^61^). Then, prior to computing the weights, the orientations were reduced (from three to one) by using the singular value decomposition algorithm on the matrix multiplication reported in equation (4). This procedure is widely adopted and used to simplify the beamforming output ^63, 64^.

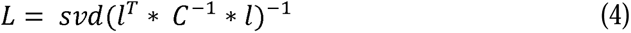

In this context, *l* denotes the leadfield model with the three orientations, while *L* is the resolved one-orientation model that was used in the estimation of the brain sources in equation (3). The weights were then applied to each brain source and timepoint, with the covariance matrix C being computed based on the continuous signal that resulted from concatenating the trials across all experimental conditions. To counterbalance the source reconstruction bias towards the head’s centre, the weights were normalised according to Luckhoo and colleagues ^64^. Since we worked on evoked responses, the weights were applied to the neural activity averaged over trials.

This procedure allowed us to obtain a time series for each of the 3559 brain sources and each experimental condition. To adjust the sign ambiguity of the evoked responses time series for each brain source, the sign was matched with the N100 response to the first tone of the auditory sequences ^26–30^.

### MEG source level and aging

For each of the significant clusters emerged from the previous analysis at the MEG sensor level, we contrasted the brain activity of young versus older adults. We averaged the time series of all brain sources over the time-window of each significant cluster and computed independent-sample t-tests contrasting the brain activity of young versus older adults. This procedure was computed independently for the three experimental conditions (M, NT1, NT3). Finally, we corrected for multiple comparisons using a 3D cluster-based MCS (α = .005 [older vs young adults], α = .05 [young vs older adults], *p*-value = .001). Here, we used a stricter α level for older vs young adults since the difference in their brain activity was particularly strong and we wanted to highlight the main focus of such differences. For this procedure, we first determined the sizes of significant clusters consisting of neighbouring brain voxels. Subsequently, we generated 1000 permutations of the initial data and estimated the sizes of significant clusters formed by neighbouring brain voxels in each permutation. This process yielded a reference distribution of the largest cluster sizes observed in the permutated data. Finally, we identified original clusters as significant if their size was larger than 99.99% of the clusters in the reference distribution. Further details on the MCS algorithm can be found in previous works by Bonetti and colleagues ^26–30^.

### Functional regions of interests (ROIs)

We computed a complementary analysis by investigating the difference between the brain activity of young versus older adults in a selected array of functional ROIs, previously described by Bonetti, Fernández Rubio, Carlomagno, Pantazis, Vuust and Kringelbach ^28^. These were derived from the whole-brain analysis of the active brain regions of young adults during recognition of the same musical sequences used in the current study. These areas roughly corresponded to the bilateral medial cingulate gyrus (MC), bilateral ventromedial prefrontal cortex (VMPFC), left (HITL) and right hippocampal area and inferior temporal cortex (HITR), and left (ACL) and right auditory cortex (ACR). In addition, we incorporated the left (IFGL) and right inferior frontal gyrus (IFGR) because these regions displayed marked differences between young and older adults.

This additional analysis allowed us to reconstruct with greater precision the time series of each brain region that played a central role in auditory sequence recognition. Thus, while it did not provide additional information to the previous analysis, it refined its significance. In **Table S4**, we reported the Montreal Neurological Institute (MNI) coordinates of each voxel forming the eight ROIs. The ROIs are visually displayed in **Figure S2**.

### Aging and ROIs time series

We contrasted the brain activity of young versus older adults by computing an independent-sample t-test for each ROI, timepoint, and condition. We corrected for multiple comparisons using 1D cluster-based MCS (α = .05, MCS *p*-value = .001). First, we identified the clusters of the significant values which were continuous in time. Second, we performed 1000 permutations, consisting of randomising the significant values obtained from the t-tests. For each permutation, we then extracted the maximum cluster size, and we built their reference distribution. To summarise, we considered significant the original clusters that were larger than the 99.99% of the clusters emerged in the permutations. Additional details on this procedure can be found in previous works by Bonetti and colleagues ^26–30^.

### WM, musical expertise, education level, aging and neural data

We computed two additional analyses to assess whether potential confounding variables had an impact on the relationship between aging and the neural responses underlying the recognition of the musical sequences.

In the first analysis we computed three independent multivariate analyses of covariance (MANCOVAs), one for each experimental condition (Wilk’s Lambda [Λ], α = .05). In each MANCOVA the dependent variables were the neural data for the eight ROIs, the independent variable was age, and the covariates were years of formal musical expertise, sex, WM, and years of formal education that participants received. To be noted, the neural data was collapsed into one single value for each ROI and participant. This was computed by averaging the main response (neural peak ± 20 ms) to each tone in the M condition. With regards to the N conditions, we selected the main response (neural peak ± 20 ms) to the tone that introduced the variation in the sequence. This analysis was conducted in R ^65^.

The second analysis consisted of computing analyses of variance (ANOVAs) for each time-point and each ROI and then using the same cluster-based 1D MCS to correct for multiple comparisons that we described in the previous paragraphs.

In this case, we computed two independent sets of ANOVAs. In the first one, we used one-way ANOVAs contrasting three age-groups: young (younger than 25), older adults 60-68 (age between 60 and 68), and older adults > 68 (older than 68). In the second set, we used two-way ANOVAs with the following levels: WM (high and low performers) and age (young and older adults). This allowed us to further test the changes in the brain activity over different age-groups as well as to better highlight the impact of WM on the ROIs time series.

**Figures 5** and **6** report the ROIs which showed the strongest results, while **Tables S6** and **S7** disclosed the complete details of the statistical results.

To be noted, four participants (three young and one older adult) did not complete the WM assessment. For this reason, the analyses described in this paragraph were computed with a sample of 72 participants.

## Data availability

The codes are available at the following links: https://github.com/leonardob92/MEG_Aging_Bach.git

https://github.com/leonardob92/LBPD-1.0.git

The multimodal neuroimaging data related to the experiment is available upon reasonable request.

## Acknowledgements

The Center for Music in the Brain (MIB) is funded by the Danish National Research Foundation (project number DNRF117).

LB is supported by Carlsberg Foundation (CF20-0239), Lundbeck Foundation (Talent Prize 2022), Center for Music in the Brain, Linacre College of the University of Oxford (Lucy Halsall fund), Society for Education and Music Psychology (SEMPRE’s 50th Anniversary Awards Scheme), and Nordic Mensa Fund.

MLK is supported by Center for Music in the Brain and Centre for Eudaimonia and Human Flourishing, which is funded by the Pettit and Carlsberg Foundations.

## Author contributions

LB, GFR, EB and MLK conceived the hypotheses. LB, GFR and ERO designed the study. LB, MLK, EB and PV recruited the resources for the experiment. LB, GFR, ERO and FC collected the data. LB, GFR and performed pre-processing and statistical analysis. MLK, EB, ML, SAK, AC and PV provided essential help to interpret and frame the results within the neuroscientific literature. GFR and LB wrote the first draft of the manuscript. LB, GFR and MLK prepared the figures. All the authors contributed to and approved the final version of the manuscript.

## Competing interests’ statement

The authors declare no competing interests.

## SUPPLEMENTARY MATERIAL

Supplementary materials related to this study and organised as supplementary figures (*i*) and tables (*ii*). In the cases when the supplementary tables were too large to be reported in the current document, they have been exported to Excel files that can be found at the following link:

https://drive.google.com/drive/folders/1mCDD1Eghm5W7aJ9jtjl-9NczNB457ROQ?usp=sharing

## SUPPLEMENTARY FIGURES

**Figure S1.**
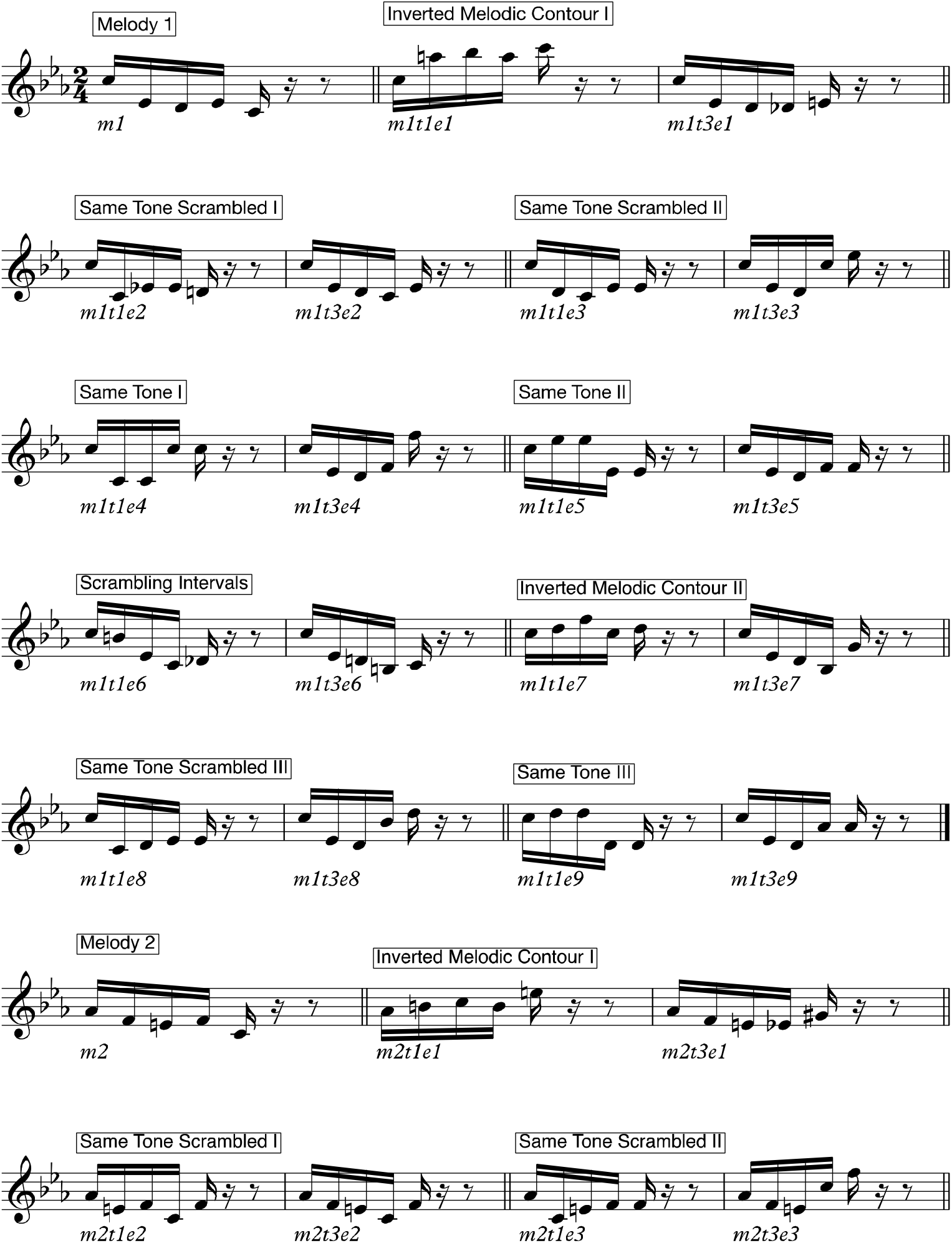
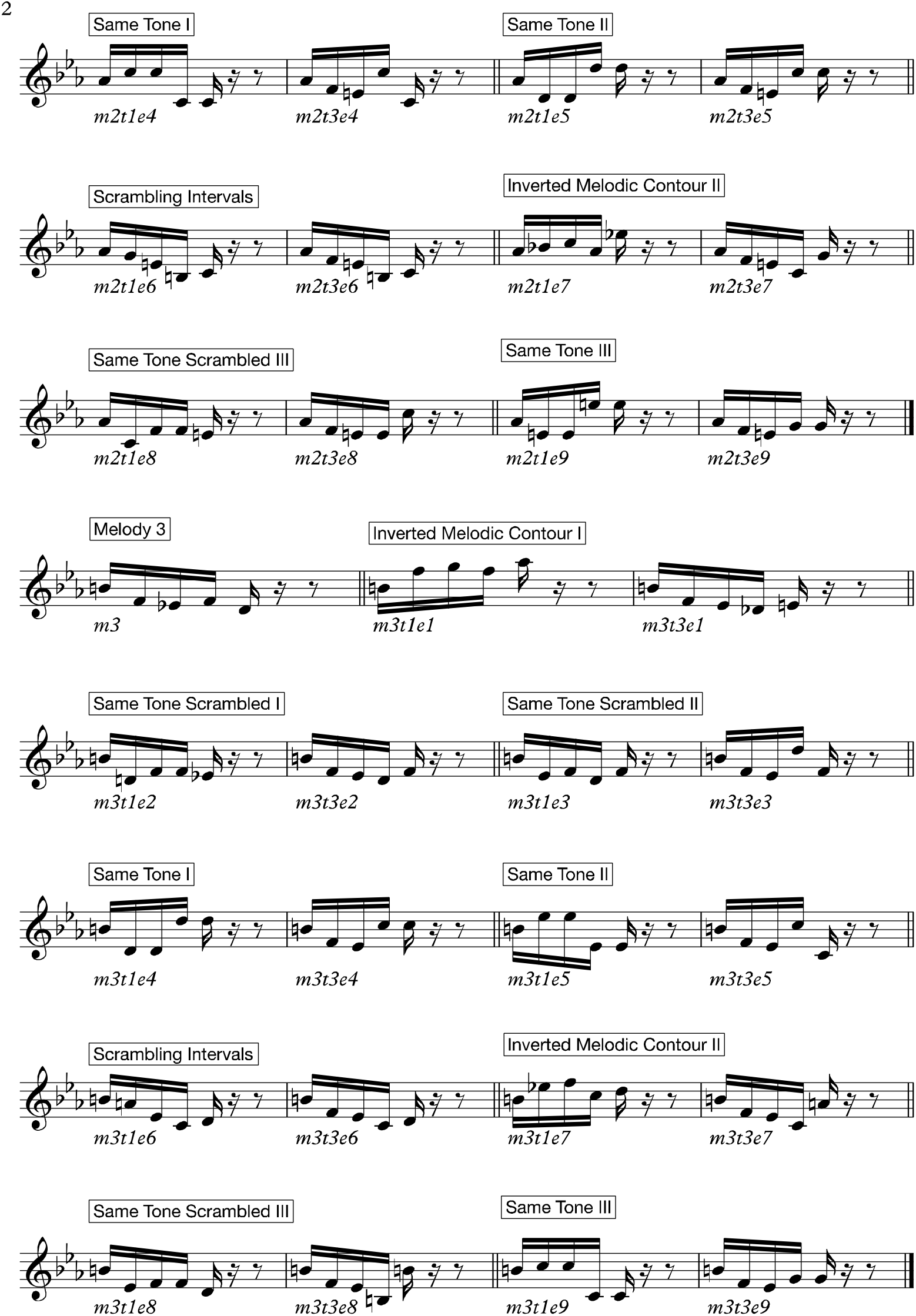
Temporal sequences used in the experiment. The figure shows all temporal sequences used in the experiment, providing detailed information on how they were created. The M sequences were three and comprised the first five tones of the first three measures of the musical piece. These three sequences were presented nine times each, for a total of 27 trials. The N sequences were created through systematic variations of the three M sequences. This procedure consisted of changing every musical tone of the sequence after the first (NT1) or third (NT3) tone. We created nine variations for each of the original M sequences and each of the four categories of N. This resulted in 27 N sequences for each category, and 54 N in total. To be noted, as shown in this figure, the variations were created according to the following rules: (i) Inverted melodic contours (used twice): the melodic contour of the variation was inverted with respect to the original M sequence (i.e., if the M sequence had the following melodic contour: down-down-up-down, the N sequence would be: up-up-down-up); (ii) Same tone scrambled (used three times): the remaining tones of the M sequence were scrambled (e.g., M sequence: C-E-D-E-C, was converted into NT1 sequence: C-C-E-E-D); (iii) Same tone (used three times): the same tone was repeatedly used, in some cases varying only the octave (e.g., M sequence: C-E-D-E-C, was transformed into NT1 sequence: C-E^8-^ E^8-^ E_8_^-^ E_8_); (iv) Scrambling intervals (used once): the intervals between the tones were scrambled (e.g., M sequence: 6^th^m - 2^nd^m – 2^nd^m – 3^rd^m, was adapted to NT1 sequence: 2^nd^m, 6^th^m, 3^rd^m, 2^nd^m).

**Figure S2.**
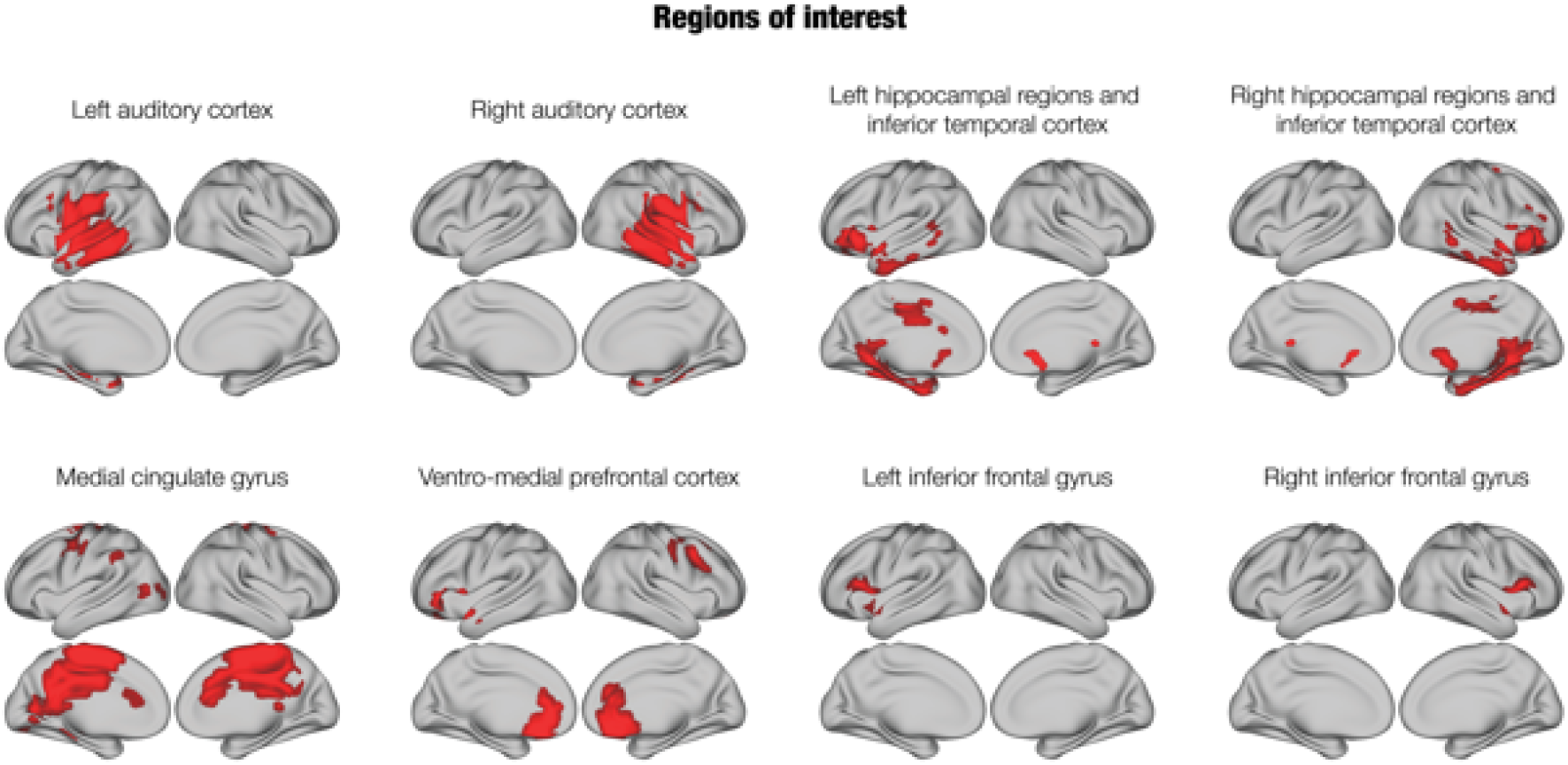
Brain parcellation. The eight ROIs used in the study: left (i) and right auditory cortex (ii); left (iii) and right hippocampal regions and inferior temporal cortex (iv); medial cingulate gyrus (v), ventromedial prefrontal cortex (vi); left (vii) and right inferior frontal gyrus (viii).

**Figure S3.**
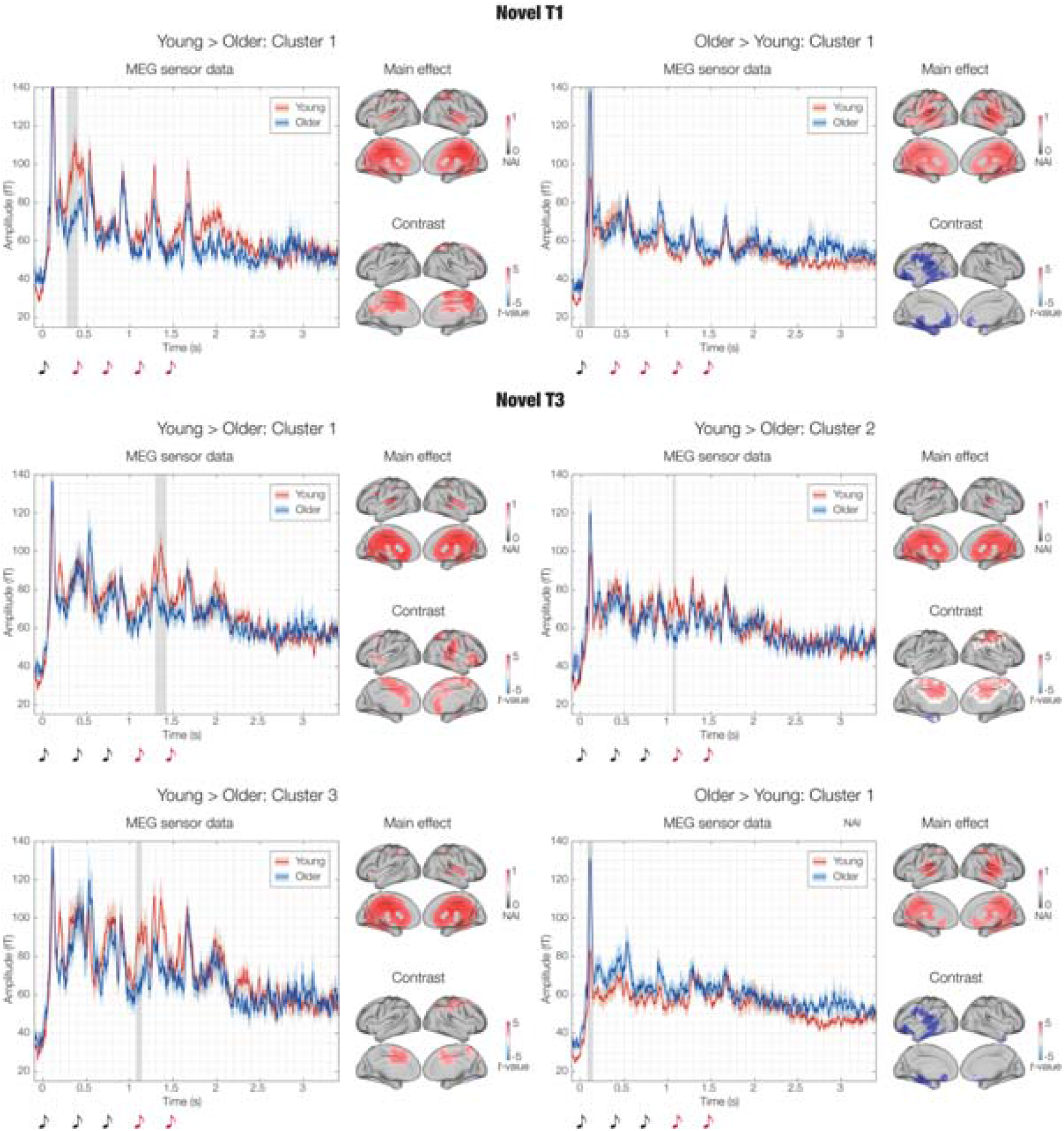
Impact of aging on the brain activity underlying the recognition of previously memorised musical sequences. Significant contrasts between the brain activity of young and older adults during the recognition of previously memorised musical sequences. For each significant cluster, the left plot shows the amplitude of the brain signal recorded for young (red) and older adults (blue). Shaded red and blue areas depict standard errors, while grey areas refer to the significant time-window for the cluster. The plot refers to the average over the absolute values of the magnetometer channels forming the significant clusters outputted by the MEG sensors MCS. The plot on the right shows the neural sources in the time-window of the significant MEG sensors cluster. The top plot shows the main effect over all participants (the colorbar indicates the reconstructed brain activity standardised between 0 and 1), while the bottom plot shows the contrast between the brain activity of young versus older adults (the colorbar indicates the t-value of the contrast). The first five clusters refer to the contrasts where the brain activity was stronger for young versus older adults. The last cluster refers to the contrasts where the brain activity was stronger for older versus young adults. ***Table 2*** reports the key statistics of these analyses, while **Table S1** shows the complete results.

**Figure S4.**
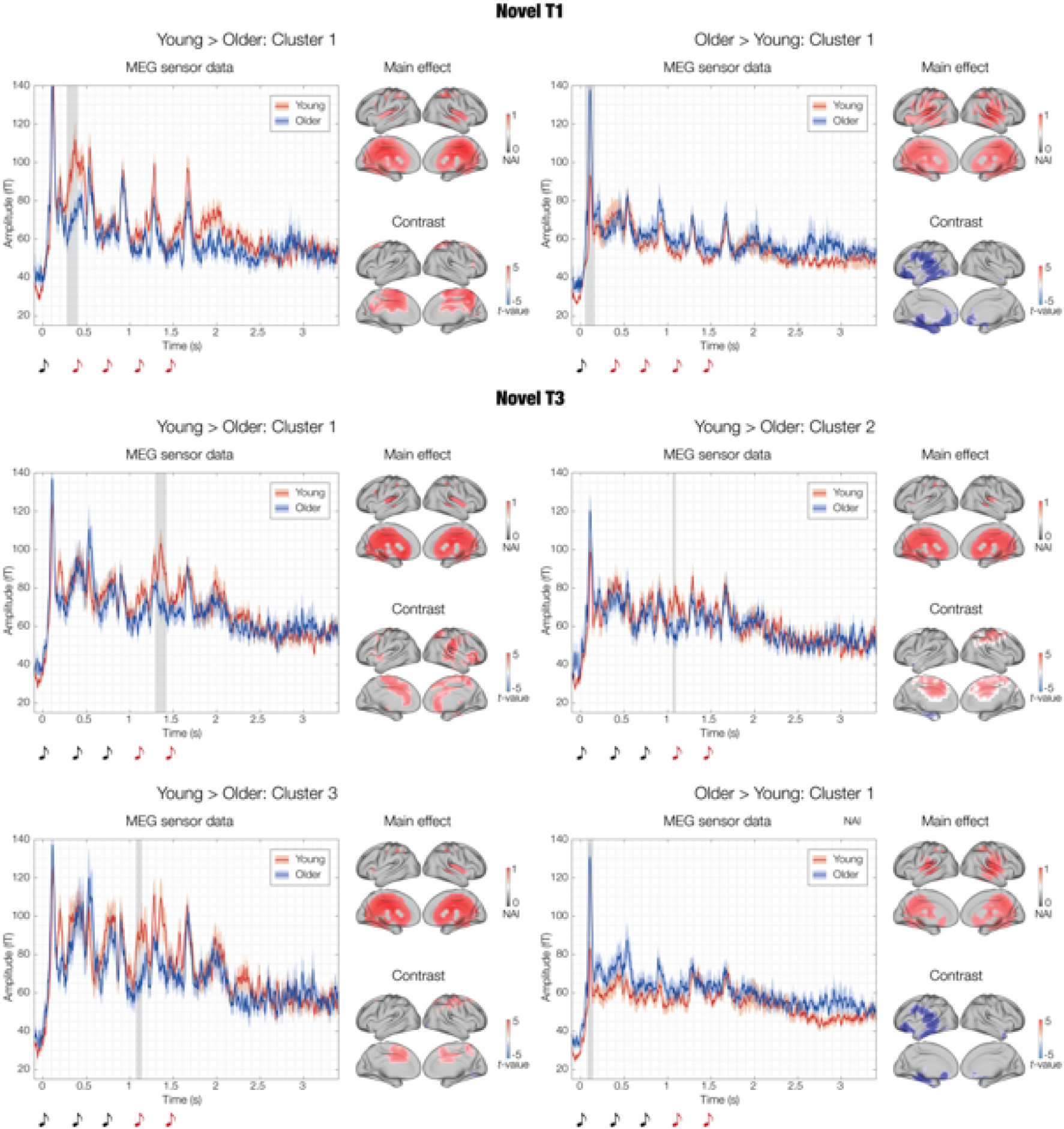
Impact of aging on the brain activity underlying the recognition of the varied (novel) musical sequences. Significant contrasts between the brain activity of young and older adults during the recognition of the varied musical sequences (NT1 and NT3). For each significant cluster, the left plot shows the amplitude of the brain signal recorded for young (red) and older adults (blue). Shaded red and blue areas depict standard errors, while grey areas refer to the significant time-window for the cluster. The plot refers to the average over the absolute values of the magnetometer channels forming the significant clusters outputted by the MEG sensors MCS. The plot on the right shows the neural sources in the time-window of the significant MEG sensors cluster. The top plot shows the main effect over all participants (the colorbar indicates the reconstructed brain activity standardised between 0 and 1), while the bottom plot shows the contrast between the brain activity of young versus older adults (the colorbar indicates the t-value of the contrast). The first two clusters refer to NT1 (on the left the contrasts where the brain activity was stronger for young versus older adults and on the right vice versa). The last four clusters refer to NT3 (the first three clusters relate to tthe contrasts where the brain activity was stronger for older versus young adults, while the last one vice versa). ***Table 2*** reports the key statistics of these analyses, while **Table S1** shows the complete results.

## SUPPLEMENTARY TABLES

***Table S1. Detailed information on significant clusters for MEG sensor data***

*Significant clusters of MEG sensors emerged from MCS contrasting the brain activity of young versus older adults, independently for the three experimental conditions (M, NT1, NT3). The table illustrates the clusters with regards to significant channels, sizes, maximum t-values and time-windows.*

***Table S2. Source reconstruction main effect.***

*Main effect for the source reconstruction performed in the time-windows of the significant clusters at MEG sensor level. Results are reported independently for each cluster and contrast, and comprise the brain region, brain hemisphere, standardised neural index and MNI coordinates for each voxel.*

***Table S3. Young versus older adults in MEG source space.***

*Significant MEG source clusters of differential brain activity between young and older adults performed in the time-windows of the significant clusters at MEG sensor level. Results are reported independently for each cluster and contrast, and comprise the brain region, brain hemisphere, t-value and MNI coordinates for each voxel.*

***Table S4. ROIs coordinates.***

*MNI coordinates for each of the voxels forming the eight ROIs.*

***Table S5. ROIs time series.***

*Significant clusters of differential brain activity between young and older adults for the eight ROIs used in the study. Results are reported independently for the eight ROIs and for each experimental condition (M, NT1, NT3), and comprise cluster size, p-value, temporal extent of the clusters and peak t-value within the cluster.*

***Table S6. ROIs time series and WM.***

*Significant clusters of differential brain activity observed by contrasting the following four categories of participants: young adults with high WM (i), young adults with low WM (ii), older adults with high WM (iii) and older adults with low WM (iv). Results are reported independently for the eight ROIs and for each experimental condition (M, NT1, NT3), and comprise cluster size, p-value, temporal extent of the clusters and peak F-value within the cluster.*

***Table S7. ROIs time series and three age groups.***

*Significant clusters of differential brain activity observed by contrasting the following three categories of participants: young adults (i), older adults aged between 60 and 68 years old (ii) and older adults older than 68 years old (iii). Results are reported independently for the eight ROIs and for each experimental condition (M, NT1, NT3), and comprise cluster size, p-value, temporal extent of the clusters and peak F-value within the cluster.*

## Notes

### Competing Interest Statement

The authors have declared no competing interest.

